# Full-thickness spatial transcriptomics of the human uterus reveals basalis niche architecture and regeneration gradients during menstrual breakdown

**DOI:** 10.64898/2026.06.08.730878

**Authors:** Valentina Lorenzi, Marie Moullet, Cecilia Icoresi-Mazzeo, Ryan Colligan, Molly R. Danks, Celeste E. Cohen, Antonio Parraga, Christina E. Kim, Carmen Sancho-Serra, Rina Sakata, Jitka Hausnerová, Karel Crha, Martina Vodičková, Andrea Hledíková, Dominika Rieglová, Martina Tomková, Josep Marí-Alexandre, Sarai Tomás-Pérez, Kristina Aghababyan, Krishnaa T. A. Mahbubani, Kourosh Saeb-Parsy, Charlotte Cassie, Tamara Garrido, Inmaculada Moreno, Aymara Mas, Felipe Vilella, Martin Prete, Nadav Yayon, Vít Weinberger, Vicente Pérez-García, Juan Gilabert-Estellés, Carlos Simon, Sarah A. Teichmann, Iva Kelava, Luz Garcia-Alonso, Roser Vento-Tormo

**Affiliations:** Wellcome Sanger Institute, Cambridge, UK; Loke Centre for Trophoblast Research, University of Cambridge, Cambridge, UK; Department of Pathology, University Hospital Brno, Brno, Czech Republic; Faculty of Medicine, Masaryk University, Brno, Czech Republic; Department of Gynecology, Obstetrics and Neonatology, University Hospital Brno, Brno, Czech Republic; Research Laboratory in Biomarkers in Reproduction, Obstetrics and Gynaecology, Research Foundation of the General University Hospital of Valencia Consortium, Valencia, Spain; Department of Pathology, General University Hospital of Valencia Consortium, Valencia, Spain; Department of Obstetrics and Gynaecology, General University Hospital of Valencia Consortium, Valencia, Spain; Department of Surgery, University of Cambridge, Cambridge, UK; NIHR Cambridge Biomedical Research Centre, Cambridge Biomedical Campus, Cambridge, UK; Carlos Simon Foundation, INCLIVA Health Research Institute, Valencia, Spain; Cambridge Stem Cell Institute, Jeffrey Cheah Biomedical Centre, Cambridge Biomedical Campus, University of Cambridge, Cambridge, UK; Centro de Biología Molecular Severo Ochoa, Madrid, Spain; Centro de Investigación Príncipe Felipe, Valencia, Spain; Department of Paediatrics, Obstetrics and Gynaecology, University of Valencia, Valencia, Spain; Department of Obstetrics and Gynecology, Beth Israel Deaconess Medical Center, Harvard Medical School, Boston, MA, USA; Department of Medicine, University of Cambridge, Cambridge, UK; CIFAR Macmillan Multi-scale Human Programme, CIFAR, Toronto, Canada

## Abstract

The human endometrium (uterine lining) undergoes cyclical breakdown and scarless regeneration during each menstrual cycle, representing an exceptional model of adult tissue renewal. Regeneration is driven primarily by progenitor cells retained within the deep, basalis compartment during menstruation, yet the full-depth spatiotemporal dynamics of this process have remained understudied due to anatomical and technical limitations. Here, we map spatial gene-expression gradients across the full thickness of the human endometrium, from the myometrial-endometrial boundary to the luminal surface, using high-resolution spatial transcriptomics integrated with single-cell transcriptomics. We profile more than ten million cells from biopsies, hysterectomy samples and menstrual fluid, enriching for the menstrual and proliferative phases, which are underrepresented in previous studies. We show that endometrial breakdown, regeneration and rapid luminal re-epithelialisation are concurrent rather than temporally separated, organised across distinct tissue compartments, revealing a mode of tissue renewal in which shedding and repair operate simultaneously. Continuous basalis-to-luminal transcriptional gradients link progenitor identity, niche signalling, and tissue remodelling, defining a coordinated regenerative axis spanning the full tissue depth. We resolve the basalis epithelial niche at unprecedented molecular resolution, identifying for the first time a discrete, predominantly quiescent progenitor-like epithelial subset and specialised supporting *SFRP5*+ fibroblasts, both characterised by WNT inhibition, alongside lymphoid aggregates, forming a multi-component architecture that persists after menopause, consistent with a long-lived regenerative reservoir. Together, these findings establish spatial transcriptional gradients as a central organising principle of endometrial renewal, providing a molecular framework for understanding disorders of menstruation, implantation failure, and impaired tissue repair.

## Main

The endometrium is the most dynamically regenerative tissue in the adult human body. Under cyclical hormonal control, the upper functionalis layer undergoes cyclical breakdown, shedding, and reconstruction, regenerating the tissue without scarring approximately 400 times across a woman’s reproductive life^1^. Disruption of this regenerative programme underlies a spectrum of common gynaecological conditions, including heavy menstrual bleeding (HMB), endometriosis and adenomyosis, in which endometrial-like cells aberrantly seed and grow outside their normal location^2^, and Asherman’s syndrome, where failure of scarless repair post injury leads to intrauterine adhesion formation and loss of endometrial function^3,4^.

Central to regeneration is the spatial organisation of the tissue along its depth. The basalis layer, retained during menstruation, is thought to harbour the progenitor epithelial populations driving post-menstrual reconstruction. Functional evidence supports an epithelial stem/progenitor compartment in the basalis: clonogenic assays established that rare epithelial progenitors with self-renewal and differentiation potential reside in the endometrium^5–7^, and subsequent studies identified candidate markers including SSEA-1^8^ and N-cadherin (*CDH2*)^9^, defining partially overlapping populations localised predominantly to basalis glands. *In vivo* lineage tracing in the mouse has provided complementary evidence for long-lived epithelial progenitors regenerating both glandular and luminal epithelium, localised either at the luminal-glandular epithelial junction^10^ or the myometrial-endometrial junction^11^. Despite functional evidence, the cellular and molecular principles organising progenitor maintenance and cyclical remodelling across the full human endometrial thickness remain incompletely understood, limiting mechanistic insight into the disorders that arise when this regenerative programme fails.

A central reason for this gap is anatomical and technical. The thin basalis layer lies at the deepest extent of the endometrium and is not reliably sampled by endometrial biopsies, which are used in the vast majority of research and clinical sampling contexts but predominantly capture the functionalis. As a consequence, the basalis has been systematically underrepresented in studies, including recent single-cell and spatial transcriptomic atlases^12–19^. Even among hysterectomy-based single-cell studies with physical access to the basalis^20–22^, individual cohort sizes and the lack of high-resolution spatial profiling have been insufficient to resolve transcriptionally subtle progenitor populations residing in this thin layer. The absence of well-defined basalis populations in existing atlases therefore reflects both anatomical sampling bias and statistical power rather than their biological absence.

Beyond the basalis, the menstrual phase itself has been relatively understudied at single-cell and spatial resolution, with studies primarily focused primarily on the window of implantation rather than the full menstrual cycle^23–25^. Progesterone withdrawal activates NF-κB signalling in decidualized stromal cells, driving cytokine and chemokine release, leukocyte recruitment, metalloproteinases upregulation, and prostaglandin synthesis^26,27^, culminating in extracellular matrix degradation and piecemeal tissue shedding^28,29^. This process embodies a striking paradox: the acute inflammatory environment required to drive tissue breakdown must be rapidly resolved to enable scarless repair^30,31^. Transient localised hypoxia^32^, glucocorticoid signalling via macrophages^33^, prostaglandin signalling and interleukin-mediated cascades^34^ have each been implicated in this resolution, yet how these signals are spatially and temporally organised across the tissue depth, which cell states execute them and how their activity is regulated in time, remains incompletely understood.

Menstrual fluid offers a complementary, non-invasive window: it provides direct transcriptomic access to the shed cellular fraction, yet its relationship to retained endometrial populations has not been systematically explored. Prior single-cell studies in humans have profiled menstrual fluid either without stage-matched eutopic comparisons or with cross-tissue comparisons restricted to stromal lineages, leaving epithelial and immune compartments uncharacterised^35^, or have examined menstrual fluid primarily in disease contexts rather than in relation to eutopic endometrial biology during menstruation and repair^36^.

The post-menopausal endometrium also remains underexplored, having been largely excluded from recent atlases focused on the reproductive years^17,21,37^. During menopause, declining ovarian estrogen drives progressive endometrial involution, producing an atrophic tissue that shares histological features with the pre-menopausal basalis^38^. Residual glands retain estrogen and progesterone receptor expression and remain capable of proliferating in response to exogenous hormone administration^38^. The few available bulk transcriptomic comparisons of pre- and postmenopausal endometrium suggest broad similarity between pre-and postmenopausal endometrium^39^, but whether cellular composition, identity and signaling are preserved at single-cell resolution remains unknown. Together, these features make the postmenopausal endometrium a unique window into the molecular organisation of the basalis, one that standard sampling approaches cannot reliably access in cycling tissue.

We set out to understand how spatial organisation across the full endometrial depth coordinates menstrual shedding and scarless regeneration, and what molecular principles define and maintain the basalis niche across the reproductive lifespan. For this, we generated a full-thickness single-cell and spatial transcriptomic atlas of the human endometrium, spanning the menstrual, proliferative and secretory phases, as well as exogenous hormone-treated conditions. By mapping cells to a continuous basalis-to-luminal axis across the full endometrial thickness, we reveal previously unreported, transcriptionally distinct basalis-restricted populations in both epithelium and stroma, establish spatial transcriptional gradients as a central organising principle of endometrial renewal, and link tissue spatial organisation to the concurrent processes of shedding and scarless regeneration. Finally, we characterise the basalis niche in postmenopausal endometrium, revealing its persistence as a long-lived regenerative compartment with immunoregulatory properties.

## Results

### A full thickness single-cell and spatial atlas of the human endometrium across the menstrual cycle

Understanding how the endometrium sheds, repairs and regenerates each cycle requires spatial resolution across the full depth of the tissue, from the myometrial boundary to the luminal surface. To resolve this organisation, we profiled 15 full-thickness hysterectomy sections spanning menstrual, proliferative, secretory, and exogenous progestogen-treated states using complementary spatial transcriptomic platforms, including 10x Genomics *Xenium* 480-plex (7,778,023 cells), *Xenium* 5k+100-plex (1,234,560 cells), and *Visium HD* (813,264 cells) (**Fig. 1a,b, Supplementary Table 1-2**). Sections were oriented from the myometrial boundary to the luminal surface, ensuring consistent inclusion of the basalis compartment across all samples – a requirement not achievable using biopsy-based spatial approaches^12–14^.

**Figure 1.**
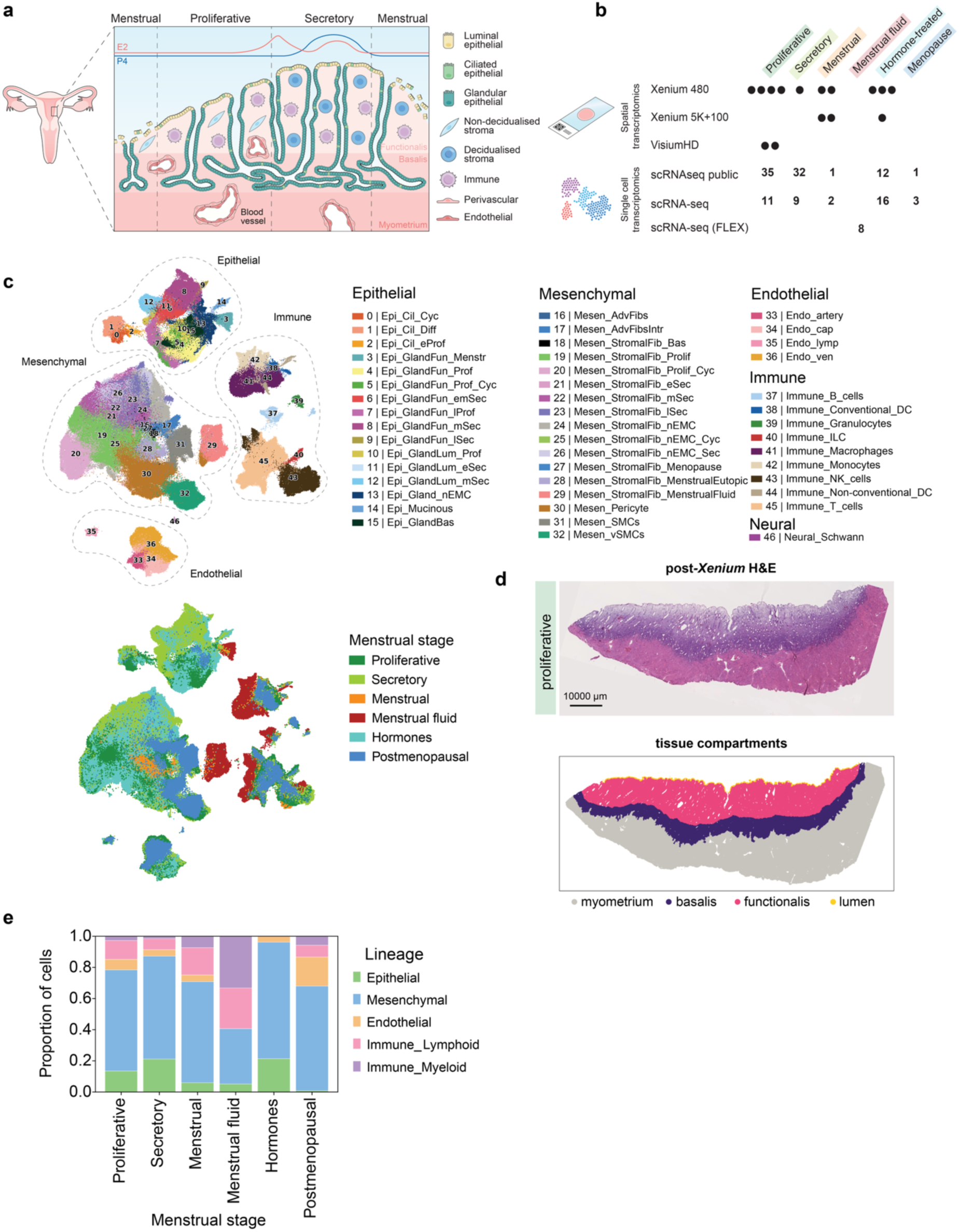
A spatially resolved single-cell atlas of the human eutopic endometrium across the menstrual cycle. a, Schematic of the human uterus showing endometrial architecture, including the functionalis and basalis layers and underlying myometrium, alongside major cell types and their distribution across menstrual cycle phases (menstrual, proliferative, and secretory). Systemic estradiol (E2) and progesterone (P4) levels are indicated above. **b**, Overview of the multimodal dataset, comprising spatial transcriptomics (*Xenium* 480-plex, *Xenium* 5K+100-plex, *Visium* HD) and single-cell RNA sequencing (newly generated scRNA-seq, scRNA-seq FLEX, and publicly available datasets) across proliferative, secretory, menstrual, menstrual fluid, hormone-treated, and postmenopausal states. Sample sizes are indicated by dots for spatial transcriptomics datasets and numbers for single-cell transcriptomics datasets. **c,** Uniform Manifold Approximation and Projection (UMAP) embeddings of uterus scRNA-seq cells in the Human Female Reproductive System Cell Atlas v1 coloured by broad cell type identity (top) and menstrual cycle stage (bottom). **d,** Representative post-*Xenium* hematoxylin and eosin (H&E) image of an endometrial tissue section in proliferative phase (top) with corresponding tissue compartment annotation (bottom), delineating myometrium, basalis, functionalis, and lumen. **e**. Bar plot showing the proportional contribution of each lineage across menstrual cycle stages.

We additionally generated 379,842 single-cell transcriptomes from 49 donors encompassing whole uteri and endometrial biopsies across the menstrual phase spectrum, 3 post-menopausal donors, and menstrual fluid collected from 8 independent donors, providing direct transcriptomic access to the shed cellular fraction during menstruation (**Fig. 1b**, **Supplementary Table 3**). Cell-state annotation was performed using the *Human Female Reproductive System Cell Atlas v1*, a harmonised reference of publicly available scRNA-seq datasets spanning multiple female reproductive tissues which integrates datasets newly generated in this study (Cohen et al., 2026, *biorxiv*), enabling consistent cell-type assignment across samples and platforms (**Fig. 1c**, **Extended Data Fig. 1a,** cell type markers reported in **Supplementary Table 4**). The scRNA-seq data for the current study is available at: https://www.reproductivecellatlas.org/HCAreproductive/v1/uterus/.

Histological images associated with each spatial transcriptomics section were reviewed by a pathologist to confirm normal endometrial architecture, and used to define tissue compartments through a two-step annotation workflow. Morphological clusters derived from the computational pathology foundation model *UNI2*^40^ first provided an automated tissue partition, which was subsequently reviewed and manually refined using *TissueTag*2^41^ (**Extended Data Fig. 1b**; Methods). Four compartments, myometrium, basalis, functionalis, and lumen, were delineated in each tissue section, with the basalis consistently represented across all samples (**Fig. 1d**, **Extended Data Figs. 1c** and **2a**).

**Figure 2.**
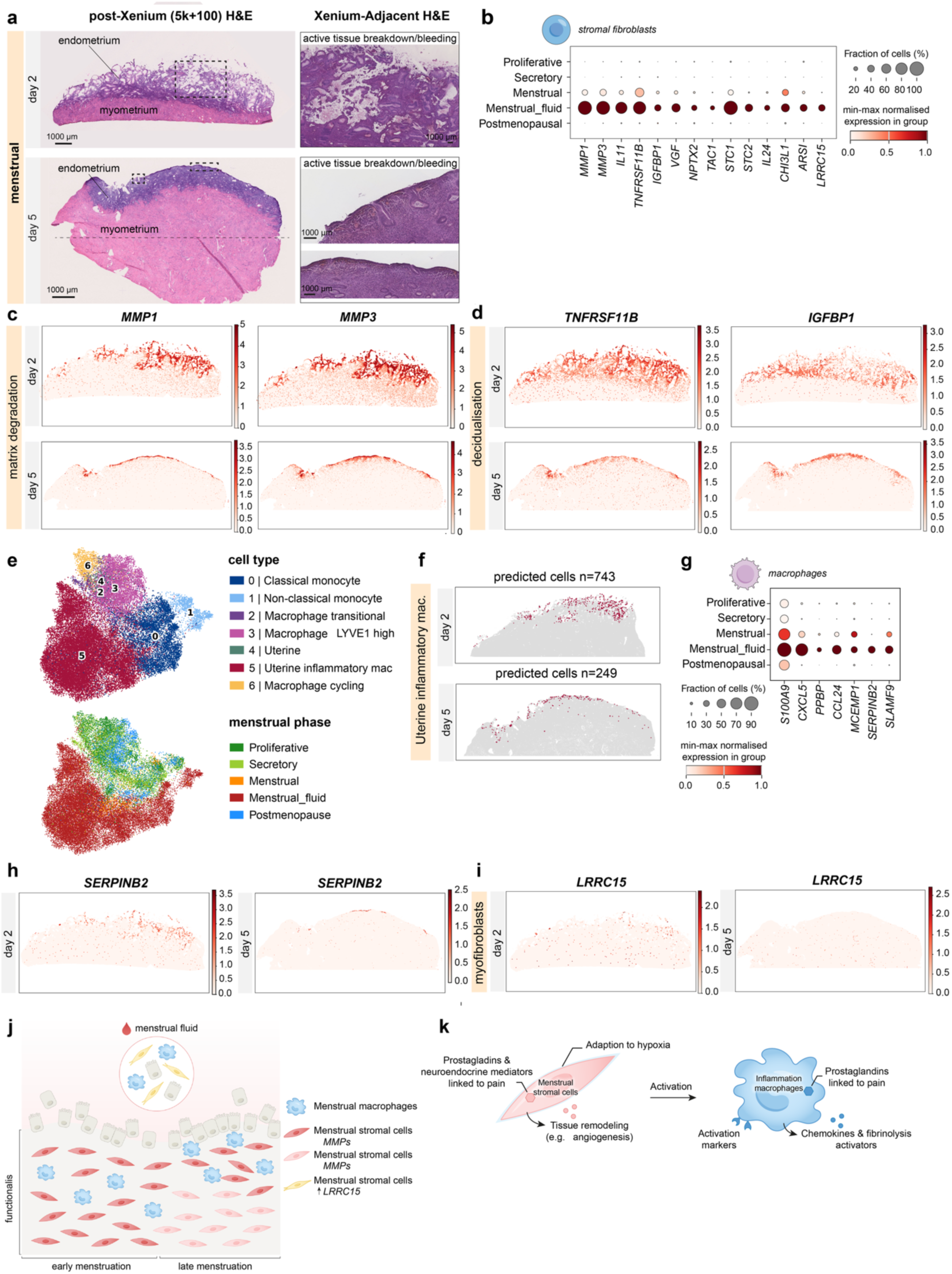
Spatially ordered stromal and macrophage programmes of breakdown and survival characterise menstrual shedding. a, Left: Post-*Xenium* (5K+100-plex) H&E images endometrial sections from menstrual days 2 (top) and 5 (bottom). The horizontal line in day 5 endometrium image indicates the lower tissue boundary shown in spatial transcriptomics analysis. Dashed rectangles highlight regions shown at higher magnification on the right. Right: H&E images of Xenium-adjacent tissue sections to the samples shown on the left, displaying active tissue shedding and bleeding. **b,** Dot plot showing expression of menstrual fibroblast markers in stromal fibroblasts across the menstrual cycle in scRNA-seq data. Dot size indicates the fraction of expressing cells and colour intensity indicates min-max normalised expression within each group. **c,** Log-normalised expression of menstrual stromal markers related to matrix breakdown in *Xenium* 5K+100-plex spatial transcriptomics sections from menstrual days 2 and 5. **d,** Log-normalised expression of menstrual stromal markers related to decidualisation in *Xenium* 5K+100-plex spatial transcriptomics sections from menstrual days 2 and 5. **e**, UMAP embedding of scRNA-seq myeloid cells, coloured by fine cell type annotation (top) and menstrual cycle stage (bottom). **f,** Inferred spatial localisation of uterine inflammatory macrophages (defined in single-cell RNA-seq) in *Xenium* 5K+100-plex spatial transcriptomics sections from menstrual days 2 and 5. **g,** Dot plot showing expression of menstrual macrophage markers across macrophage populations throughout the menstrual cycle in scRNA-seq data. **h,** Log-normalised expression of the macrophage-derived antifibrinolytic factor *SERPINB2* in *Xenium* 5K+100-plex spatial transcriptomics sections from menstrual days 2 and 5. **i,** Log-normalised expression of menstrual stromal markers related to myofibroblast activation in *Xenium* 5K+100-plex spatial transcriptomics sections from menstrual days 2 and 5. **j,** Schematic illustration of the proposed model of stromal and myeloid dynamics during menstruation. **k,** Schematic illustration summarising proposed functions of menstrual stromal cells and macrophages.

Cell states were then mapped onto the spatial transcriptomic datasets using two computational approaches: ISS-patcher^42^, which is kNN-based, and TACCO^43^, which is optimal transport-based. TACCO achieved the best balance between aggregate concordance between single-cell and spatial expression profiles of the same predicted cell types and accurate mapping of rare cell types, and was therefore used for all downstream analyses (**Supplementary Note, Extended Data Fig. 2b-d**). Integrating spatial cell-state mapping with morphological compartment annotations resolved the distribution of endometrial cell states across tissue depth and cycle phases, revealing spatially restricted populations characterised in subsequent tissue sections. The number of cell types predicted in each compartment per stage (summed across donors) are reported in **Supplementary Table 5.**

Understanding the full process of menstrual breakdown requires characterising not only the retained tissue but also the shed cellular fraction. Menstrual fluid is inaccessible to spatial profiling of eutopic tissue, and thus offers a complementary window onto the cellular states active at the shedding front. Characterisation of the menstrual fluid cellular composition revealed endometrial epithelial (5% of all menstrual fluid cells), mesenchymal (35%), and immune cell types (33% myeloid and 26% lymphoid), with a notable absence of endothelial cells (<1%) (**Fig. 1e, Extended Data Fig 2e**). Epithelial and mesenchymal cells were confirmed as endometrial in origin by canonical marker expression, and shed stromal fibroblasts retained expression of decidualization markers *DKK1* and *IGFBP1* (**Extended Data Fig. 2f**), and mapped predominantly to the functionalis compartment, as expected for the shed tissue (**Extended Data Fig 2g**).

Together, these data establish a spatially and transcriptomically resolved map of the human endometrium at unprecedented depth and temporal resolution, integrating full-thickness spatial transcriptomics with single-cell profiling across cycle phases including the menstrual phase. The following sections characterise how the spatial organisation of the tissue underlies endometrial shedding, scarless regeneration and basalis niche maintenance.

### Spatially ordered stromal and macrophage programmes of breakdown and survival characterise menstrual shedding

Menstruation involves cyclical tissue breakdown triggered by progesterone withdrawal, followed by inflammatory remodelling and scarless repair. To identify the cell states expressing the molecular mediators associated with menstrual shedding and repair, we first identified genes selectively enriched during the menstrual phase (11 donors) within macrophage and stromal populations, recognised orchestrators of regenerative tissue responses^31,44,45^. We then mapped these menstrual phase-specific genes onto 2 whole-uterine menstruating samples (cycle day 2 and 5) profiled using *Xenium* spatial transcriptomics (5k+100-plex panel), enabling direct spatial characterisation of active tissue breakdown and regeneration (**Fig 2a**).

The spatial distribution of core menstrual stromal markers involved in matrix degradation (*MMP1*, *MMP3*) and decidualisation (*TNFRSF11B, IGFBP1, IL11*, *IL24*)^44,46^ revealed a strikingly ordered wave of stromal tissue breakdown and reconstruction not previously appreciated at this spatial resolution **(Fig. 2b-d**, **Extended Data Fig. 3a)**. At menstrual day 2, *MMP1/MMP3//TNFRSF11B/IGFBP1/IL11/IL24*-positive stroma occupied most of the functionalis, whereas by day 5 this programme became restricted to a thin subluminal layer, consistent with progressive resolution of the breakdown front and reconstruction of the endometrium, and closely matching the spatial distribution of bleeding and tissue disruption observed in corresponding histology sections (**Fig. 2a-d**, **Extended Data Fig. 3a**).

Following the same spatial distribution as MMP-expressing stroma, we identified a specialised inflammatory macrophage population of monocyte origin (*S100A9*+), present in both eutopic menstrual tissue and menstrual fluid (**Fig. 2e-g**). Menstrual macrophages specifically expressed chemokines involved in granulocyte recruitment, including *CXCL5* and *PPBP*, two potent neutrophil-recruiting factors linked to inflammatory amplification and prostaglandin-associated uterine contractility, as well as *CCL24*, which has been associated with eosinophil recruitment and pro-fibrotic macrophage–fibroblast signalling^47,48^ (**Fig. 2g**). They additionally upregulated *MCEMP1*, *SERPINB2*, and *SLAMF9*, consistent with activation of monocyte-derived inflammatory macrophage states associated with enhanced response to infection, migration, survival, and cytokine production^49,50^ (**Fig. 2g**). Notably, *SERPINB2* also inhibits fibrinolysis, enhancing coagulation^51^, and was spatially localised to the endometrial surface following the same pattern as inflammatory macrophages (**Fig. 2h**).

In contrast to this spatially restricted inflammatory programme, a second set of menstrual stromal markers was broadly distributed across the tissue at both time points, consistent with roles in cell survival and tissue homeostasis. This included *STC2*, a direct HIF1α target promoting survival during hypoxia and endoplasmic reticulum stress^52^ (**Fig. 2b**, **Extended Data Fig. 3b**), which may also support local tissue homeostasis by promoting an anti-inflammatory microenvironment, as its knockdown in mesenchymal stem cells reduced IL10 secretion and impaired macrophage polarisation toward a pro-repair phenotype^53^. Similarly broadly distributed was *CHI3L1,* a TGF-β induced chitinase-like glycoprotein secreted by activated macrophages and fibroblasts that promotes angiogenesis, M2 macrophage activation, and wound healing^54^ (**Fig. 2b**, **Extended Data Fig. 3c**). Interestingly, only menstrual fluid stromal cells (but not eutopic menstrual stromal cells) additionally expressed *LRRC15*, a marker of active TGF-β-activated myofibroblasts that suppresses CD8+ T cell function in the tumour microenvironment^55^, indicating that activated, pro-fibrotic stromal states were confined to the shed compartment (**Fig. 2b,i**).

Menstrual stromal cells additionally upregulated mediators linked to nociceptive signalling. Expression of *VGF*, which was spatially distributed across the endometrium at both menstrual days, together with *NPTX2* and *TAC1*, detected at the single-cell transcriptomics level, suggests previously unappreciated mechanisms contributing to menstrual pain^56–58^ (**Fig. 2b**, **Extended Data Fig. 3d**). Coordinated expression of the prostaglandin synthesis components *PTGES1*, *PTGES2*, and *PTGES3* across both stromal and macrophage populations during menstruation further pointed to the convergent prostaglandin-mediated uterine contractility and pain signalling between these two lineages during menstruation **(Extended Data Fig. 3e).**

Together, these findings reveal two spatially distinct programmes operating concurrently during menstruation: a spatially restricted inflammatory and matrix-degrading programme that retreats progressively from the functionalis by day 5 and a broader stromal programme associated with tissue survival and homeostasis. Pro-fibrotic stromal states are confined to the shed fraction, preserving the anti-fibrotic environment of the retained endometrium and consistent with the scarless character of physiological endometrial repair **(Fig. 2j-k).**

### Transient barrier remodelling and luminal identity acquisition characterise the menstrual epithelium

We next examined how the epithelial lineage contributes to the distinctive shedding and regenerative properties of the endometrium. Although the endometrium is lined by a simple columnar epithelium, we found that menstrual epithelial cells activated a transcriptional programme characteristic of stratified epithelia, encompassing terminal differentiation and barrier remodelling genes not previously associated with endometrial biology, and which was absent outside the menstrual phase. This programme included *MAB21L4*, *SCEL*, *SERPINB7*, and *PADI1*, genes more commonly associated with stratified epithelia^59–62^ (**Fig. 3a**). At menstrual day 2, *SERPINB7* spatial localisation was restricted to the uppermost portion of the shedding layer, whereas at day 5 it coincided with the sub-luminal region of tissue breakdown, consistent with progressive confinement of the barrier remodelling programme to the active shedding front (**Fig. 3b**).

**Figure 3.**
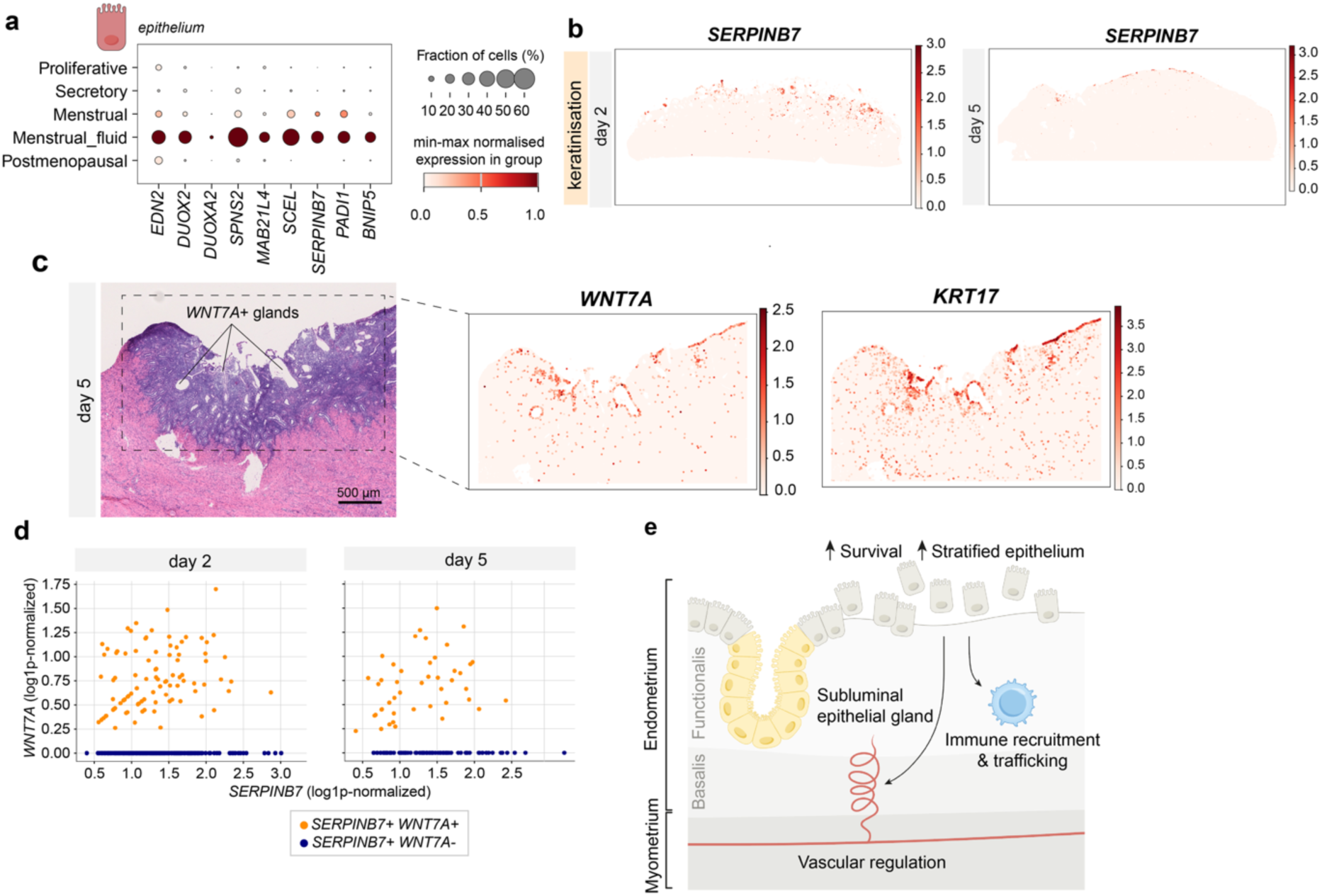
**Epithelial plasticity drives barrier remodelling and luminal repair during menstruation**. **a,** Dot plot showing expression of epithelial menstrual markers across epithelial cells throughout the menstrual cycle in scRNA-seq data. Dot size indicates the fraction of expressing cells, and colour intensity reflects min–max normalised expression within each group. **b,** Log-normalised expression of the epithelial-derived keratinisation gene *SERPINB7* in *Xenium* 5K+100-plex spatial transcriptomics sections from menstrual days 2 and 5. **c**, Left: Post-*Xenium* H&E image of a menstrual day 5 section with *WNT7A*⁺ glands annotated. Right: Corresponding log-normalised spatial expression of luminal epithelial markers *WNT7A* and *KRT17*. **d,** Scatterplot showing the expression of *WNT7A* and *SERPINB7* in epithelial spatial cells from menstruation day 2 (left) and day 5 (right). **e,** Schematic illustration showing the proposed model of epithelial cell states during menstruation and regeneration.

Shed menstrual epithelial cells also expressed *BNIP5*, a selective inhibitor of Bak-dependent apoptosis, suggesting a mechanism supporting epithelial survival during detachment from the basement membrane^63^ (**Fig. 3a**). Enhanced survival of shed epithelial cells is in keeping with their capacity to produce factors that may contribute to tissue regeneration^64^, which we explore further in our dataset. Menstrual epithelial cells in both the shed and retained fraction prominently expressed *EDN2*, a vasoconstrictive peptide that is transiently produced by ovarian granulosa cells to drive follicle rupture through smooth muscle contraction^65^, together with *DUOX2* and *DUOXA2*, which encode the epithelial NADPH oxidase complex responsible for generating extracellular hydrogen peroxide gradients that recruit leukocytes to wounded epithelia^66,67^ (**Fig. 3a**). Notably, *DUOX2* and *DUOXA2* were recently found to be induced during the post-breakdown phase of an *in vitro* menstruation model using endometrial organoids^68^ but their expression *in vivo* had not been shown to our knowledge. *In situ*, *DUOX2* expression was broadly distributed across the luminal and glandular epithelium at both menstrual day 2 and day 5, suggesting activation in both shed and retained epithelial cells (**Extended Data Fig. 3f**). Menstrual epithelial cells also expressed *SPNS2*, a transporter required to establish extracellular sphingosine-1-phosphate (S1P) gradients controlling lymphocyte and NK-cell trafficking^69^ (**Fig. 3a**).

Rapid re-establishment of a continuous luminal epithelial surface (luminal re-epithelialisation) is thought to restrict luminal immune cell access to denuded stromal tissue, limiting the inflammatory and pro-fibrotic signalling that prolonged surface epithelial absence would otherwise permit^29^. Rapid luminal re-epithelialisation through the expansion of a luminal progenitor has previously been proposed as a dominant repair mechanism in the mouse uterus^70^ and loss of *WNT7A* has recently been shown to impair long-term maintenance of endometrial organoids^68^. Here, at menstrual day 5, we found that subluminal glands expressed *WNT7A* and *KRT17*, markers that will be restricted to the lumen later on^22^, raising the possibility that luminal repair involves not only migration of residual luminal epithelium around shedding foci, but also acquisition of luminal identity by regenerating glandular epithelial cells (**Fig. 3c**). Approximately 30% of *WNT7A* cells co-expressed *SERPINB7*+, suggesting that luminal epithelial cells simultaneously adopt a transient stratified-like barrier programme during repair (**Fig. 3d**). These observations suggest a remarkable plasticity of luminal and glandular epithelial identity during repair.

Together, these findings reveal that menstrual epithelial cells are transcriptionally active participants in shedding and repair, expressing programmes associated with survival, immune-recruiting and regenerative factors, alongside evidence of luminal-glandular epithelial plasticity, collectively consistent with an active epithelial contribution to scar-free endometrial regeneration (**Fig. 3e**).

### Continuous basalis-to-luminal transcriptional gradients define the regenerative axis of the endometrium

The analyses above identified transcriptional programmes specifically expressed during menstruation, focusing on genes that are switched on or off between hormonal cycle phases. While powerful for identifying phase-specific cell states, this approach cannot detect genes whose expression persists across phases but whose spatial distribution is dynamically reorganised during each phase. To map this architecture, we constructed a continuous positional axis spanning the full endometrial depth, from the myometrial-endometrial boundary to the luminal surface, and transferred axis coordinates to the dissociated single-cell reference atlas (**Fig. 4a**, Methods). The axis was partitioned into six spatial bins reflecting the approximate proportional depth of each compartment, enabling pseudobulk generalised linear modelling of expression trends along the depth axis while reducing noise from sparse single-cell measurements and avoiding inflated false discovery rate^71^. (**Supplementary Note, Extended Data Fig. 4a-c**). TACCO-based transfer yielded higher concordance between mean scaled gene expression profiles of matched spatial and single-cell compartments than ISS-patcher-based transfer on average, and was therefore used for all subsequent analyses. Generalised linear modelling identified spatially organised genes in the epithelium and in stromal fibroblasts, independently, and found that the majority of them showed significant axis–phase interactions, indicating that spatially organised transcription is both widespread and dynamically remodelled across the menstrual cycle (**Supplementary Table 6**).

**Figure 4:**
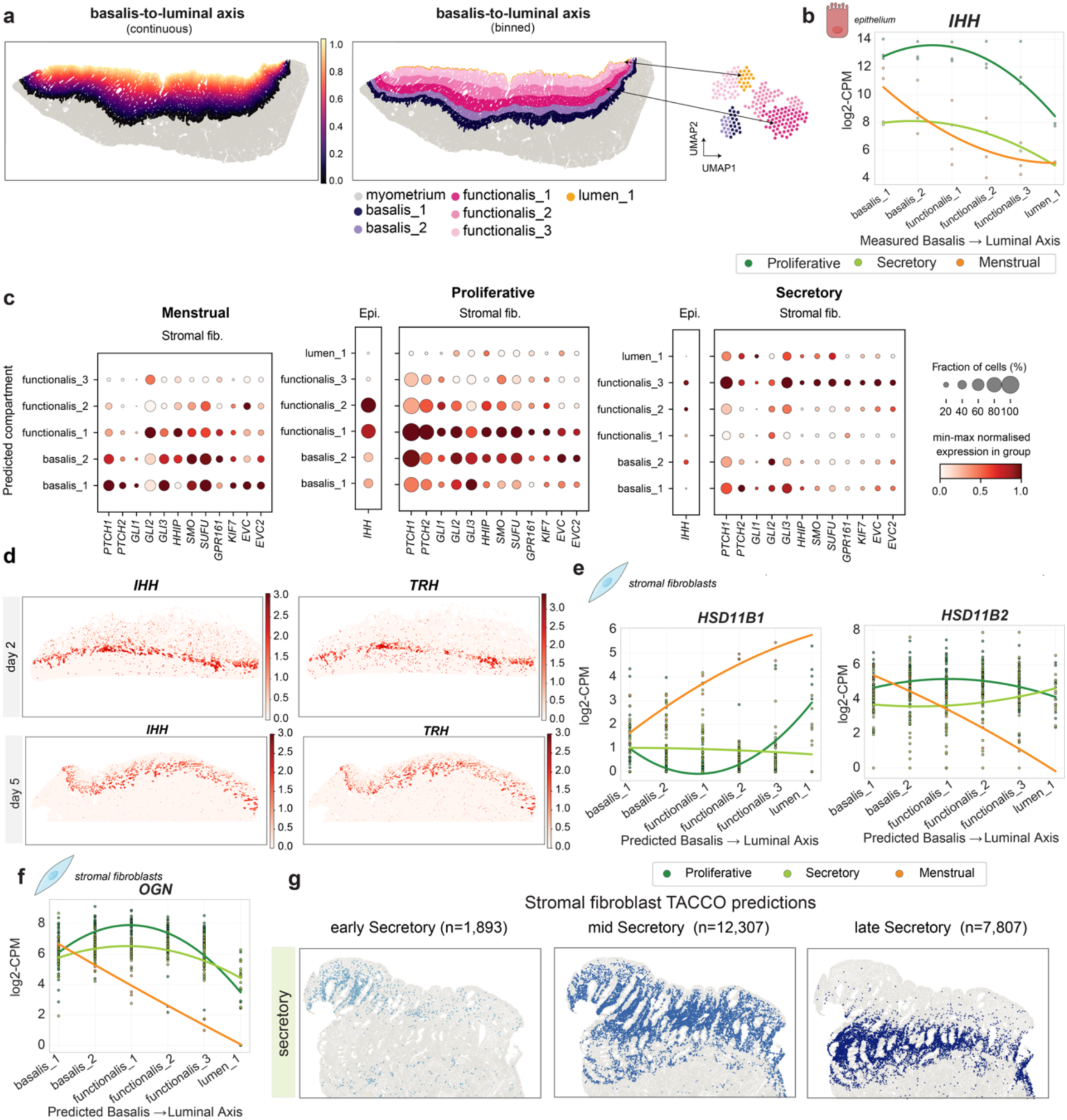
Continuous basalis-to-luminal transcriptional gradients define the regenerative axis of the endometrium. **a**, Schematic representation of the binning strategy for the basalis-to-luminal continuous axis and transfer of spatial compartment labels from spatial transcriptomics (*Xenium* 480-plex) to dissociated single cell RNA-seq data. **b**, Spline plots showing the log-normalised expression of *IHH* along the measured basalis-to-luminal binned axis in *Xenium* 480-plex spatial transcriptomics data. **c**, Dot plot showing min-max normalised expression of hedgehog ligands in epithelial cells and hedgehog target genes in stromal fibroblast cells, stratified by menstrual phase and predicted spatial compartment. Menstrual epithelial cells were not modelled due to limited epithelial representation in scRNA-seq. No fibroblasts were confidently assigned to the menstrual luminal compartment. **d**, Log-normalised expression of the epithelial-derived genes *IHH* and *TRH* in *Xenium* 5K+100-plex spatial transcriptomics sections from menstrual days 2 and 5. **e**, Spline plots showing the log-normalised expression of hydroxysteroid dehydrogenase enzymes (*HSD11B1*, *HSD11B2*) along the predicted basalis-to-luminal binned axis in scRNA-seq data. Lines are coloured by menstrual phase, and each point represents pseudobulked cells assigned to a specific bin per donor. **f**, Spline plots showing the log-normalised expression of *OGN* along the predicted basalis-to-luminal binned axis in single-cell RNA-seq data. **g**, Inferred spatial localisation of secretory stromal fibroblasts (defined by single-cell RNA-seq) in a *Xenium* 480-plex spatial transcriptomics section from secretory phase. Blue indicates cells assigned to early (left), mid (centre), or late (right) secretory fibroblast states.

Systematic ranking of genes by their epithelial basalis-to-lumen dynamic range identified *TRH*, encoding thyrotropin-releasing hormone and previously reported to be enriched in the epithelial basalis compartment^21,72^, and *IHH* encoding Indian Hedgehog, a ligand of the Hedgehog (HH) signalling pathway with an established role in endometrial regeneration^21,22,73,74^, as top candidates (**Supplementary Table 6**). Across all cycle phases, *IHH* was expressed predominantly in glandular epithelium, highest in the basalis and declining progressively toward the lumen (**Fig. 4b,c**). In stromal fibroblasts, HH pathway components, including the receptor *PTCH1*, transcriptional effectors *GLI1-3*, and modulators S*MO, SUFU, GPR161, KIF7, EVC, EVC2,* and *HHIP,* displayed a concordant spatial gradient, consistent with spatially coordinated epithelial-stromal HH signalling (**Fig. 4c**). Modelling of *GLI1* and *PTCH1* gradients in Xenium confirmed this trend (**Extended Data Fig. 4d**). Pathway activity was markedly attenuated in the secretory phase, consistent with progesterone-mediated suppression^73^ (**Fig. 4c**).

Given the pronounced spatial dynamics of the menstrual phase identified above, we examined *IHH* and *TRH* expression at menstrual days 2 and 5. At menstrual day 2, *IHH* and *TRH* were confined to the basalis (**Fig. 4d**). By day 5, both had re-expanded through the regenerating glandular epithelium (**Fig. 4d**), adjacent to residual luminal cells expressing the terminal differentiation programme described in (**Fig. 3c**). The presence of *IHH* and *TRH* expression beyond the basalis as early as menstrual day 5 indicates that regeneration is initiated while shedding is still ongoing. The progressive re-establishment of basalis-associated epithelial identity across the tissue thickness, while shedding is still ongoing, is consistent with a temporal overlap critical for preserving basalis niche integrity and preventing establishment of a fibrosis-permissive microenvironment.

We next identified genes with menstrual phase-specific spatial patterning distinct from the *IHH/TRH* axis. *HSD11B1* and *HSD11B2*, which encode the interconverting enzymes 11β-hydroxysteroid dehydrogenase types 1 and 2 by stromal cells, together regulate the local balance between inactive cortisone and its active form cortisol, a glucocorticoid with potent anti-inflammatory properties. *HSD11B1* expression is known to be induced during menstruation by inflammatory mediators including IL1^30,31^, yet its spatial organisation across the tissue has not been described. Here, we find that *HSD11B1* and *HSD11B2* display opposing gradients along the basalis-to-luminal axis. In the proliferative and secretory stages, both genes show a small dynamic range with minimal change across this axis (**Fig. 4e**). This spatial pattern is specifically reconfigured during menstruation, with the lumen showing higher *HSD11B1* and lower *HSD11B2* (**Fig. 4e).** This spatial rebalancing could generate a local gradient of increased cortisol in the lumen, providing a regionally controlled mechanism by which inflammation becomes self-limiting during tissue breakdown. Other genes showing phase-specific spatial patterning include the extracellular matrix protein osteoglycin (*OGN*), which displayed a marked decrease along the basalis-to-lumen axis specifically during menstruation (**Fig. 4f**). Previous work has established the role of *OGN* in driving fibrosis in the lung^75^, its selective downregulation during menstruation may therefore serve a protective role, limiting fibrotic remodelling during tissue breakdown.

We next examined whether variation in cellular composition within a lineage contributes to the observed gradient dynamics. In previous work, we characterised fibroblast subpopulations enriched at distinct stages of the menstrual cycle^21^, but whether these co-exist in a spatially organised manner within individual donors was unresolved. Spatial mapping of early, mid and late secretory fibroblast states onto a secretory stage sample revealed that these transcriptionally distinct populations were arranged in a progressive gradient through the functionalis. Early states occupying more luminal positions and late states localised closer to the basalis (**Fig. 4g)**. The co-occurrence of these transcriptionally distinct states within an individual donor indicates that the observed gradients reflect spatial organisation rather than inter-individual variation, and that stromal fibroblast differentiation is coordinated along both spatial and temporal axes simultaneously.

Together, these findings establish spatial transcriptional gradients as a pervasive organising principle of the endometrium, identifying HH signalling as a spatially organised, cycle-regulated epithelial-to-stromal axis, and implicate its basalis-to-lumen gradient as a coordinator of both homeostatic remodelling and regenerative re-establishment following breakdown.

### Quiescent epithelial progenitor-like cells in the human endometrial basalis

A major limitation of previous endometrial transcriptomic studies has been the inability to directly profile the basalis, the deep-lying and thin endometrial compartment retained during menstruation and thought to harbour the progenitor populations responsible for cyclical regeneration. Prior studies identified *CDH2* and *AXIN2* as enriched in basalis epithelial cells with progenitor properties^9,21,72^, yet our spatial profiling across the full endometrial thickness revealed that expression of both *CDH2* and *AXIN2* extends beyond the deepest basalis glands and is more broadly distributed across the glandular epithelium and the whole surrounding stroma than previously appreciated, undermining their utility as transcriptomic markers of the basalis (**Fig. 6a-c**).

**Figure 5.**
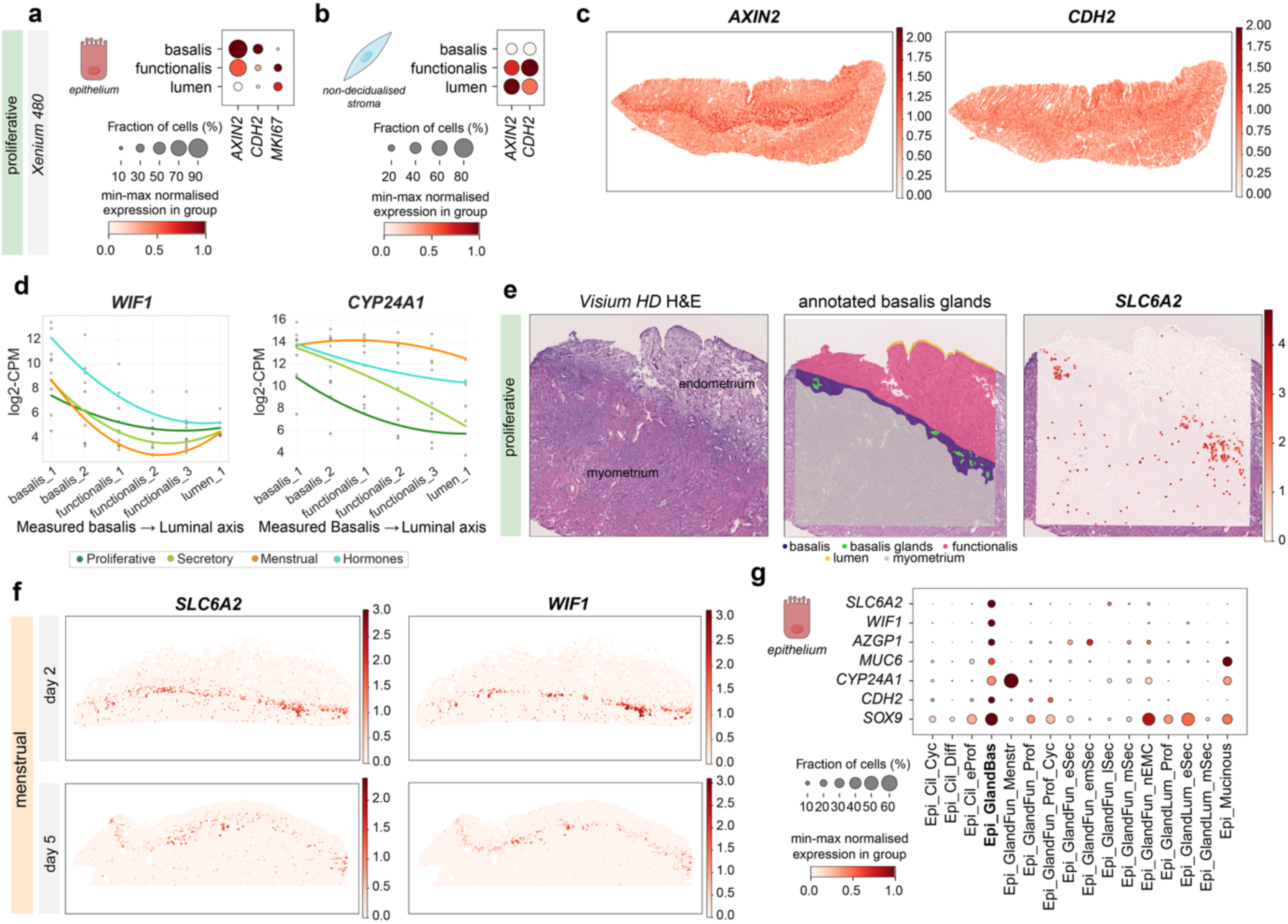
Quiescent epithelial progenitor-like cells in the human endometrial basalis. a, Dot plot showing the expression of widely reported basalis markers *AXIN2* and *CDH2* alongside cell cycle gene *MKI67* in epithelial cells in a *Xenium* 480-plex spatial transcriptomics section from proliferative phase. Dot size indicates the fraction of expressing cells, and colour intensity reflects min-max normalised expression within each group. **b,** Dot plot showing the expression of widely reported basalis markers *AXIN2* and *CDH2* alongside cell cycle gene *MKI67* in stromal fibroblast cells in a *Xenium* 480-plex spatial transcriptomics section from proliferative phase. Dot size indicates the fraction of expressing cells, and colour intensity reflects min-max normalised expression within each group. **c,** Log-normalised expression of widely reported basalis markers *AXIN2* and *CDH2* in a *Xenium* 480-plex spatial transcriptomics section from proliferative phase. **d,** Spline plots showing the log-transformed expression of *WIF1* and *CYP24A1* in epithelial cells across the basalis-to-luminal binned axis in proliferative, secretory, menstrual, and hormone-treated conditions, as measured by *Xenium* 480 spatial transcriptomics. Lines are coloured by menstrual phase, and each point represents pseudobulked cells assigned to a specific bin per donor. **e,** *Visium HD* H&E image of a proliferative endometrial section (left), with manual annotation of basalis glands (centre) and corresponding log-normalised spatial expression of *SLC6A2* (right) **f,** Log-normalised expression of the epithelial basalis genes *SLC6A2* and *WIF1* in *Xenium* 5K+100-plex spatial transcriptomics sections from menstrual days 2 and 5. **g**, Dot plot showing the expression of basalis gland epithelial markers across epithelial cell subtypes. Dot size indicates the fraction of expressing cells, and colour intensity reflects min-max normalised expression within each group.

**Figure 6:**
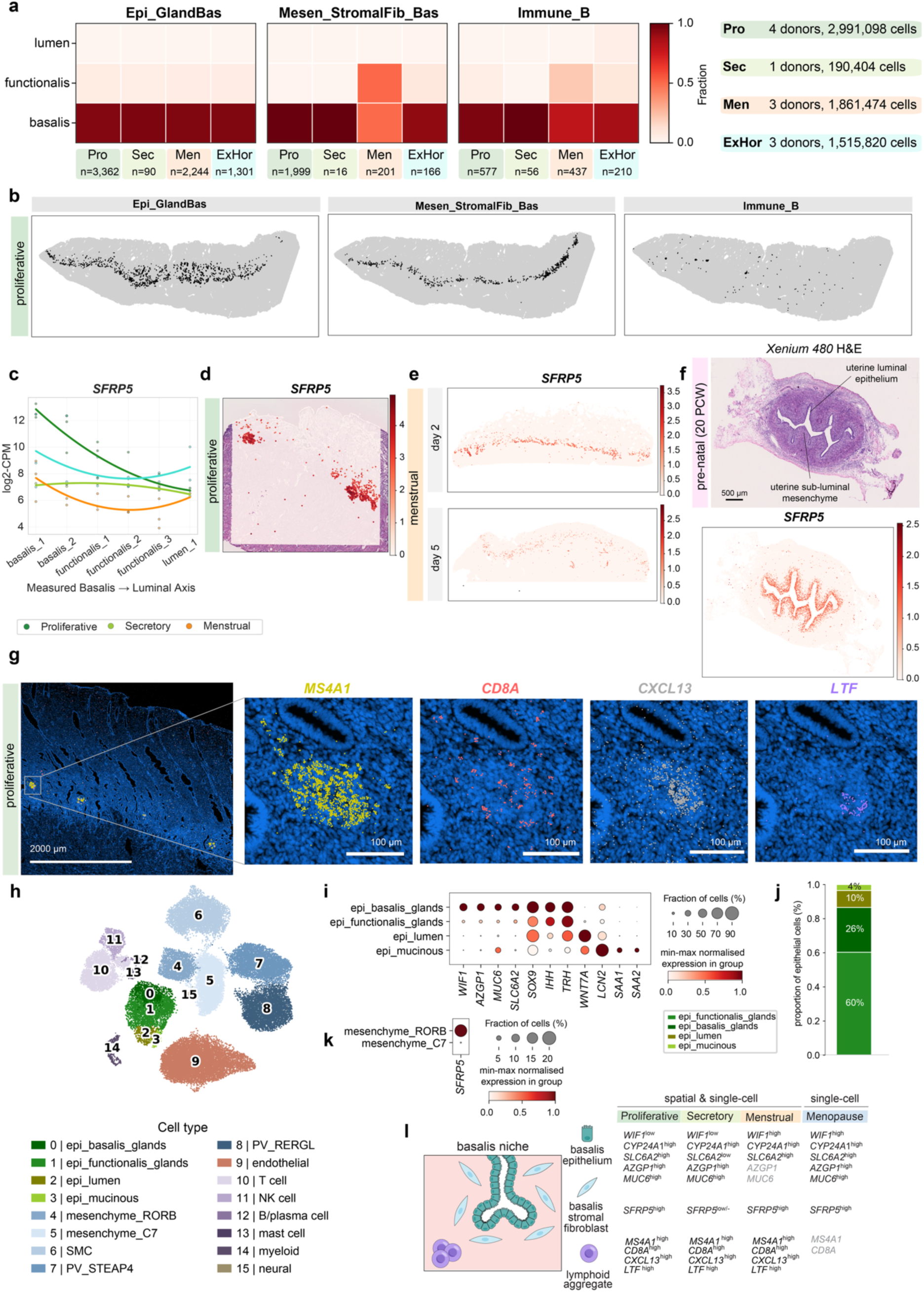
A persistent Wnt-inhibitory stromal and immune niche surrounds quiescent basalis epithelium. a, Heatmaps showing the fractional abundance of cell populations enriched in the basalis as inferred by TACCO, separated by menstrual stage. **b,** TACCO predicted spatial localisations of the cell populations enriched in the basalis reported in **a,** in a representative *Xenium* 480-plex tissue section from proliferative phase. **c,** Spline plots showing the log-transformed expression of *SFRP5* in stromal fibroblast cells across the basalis-to-luminal binned axis in proliferative, secretory, menstrual, and hormone-treated conditions, as measured by *Xenium* 480 spatial transcriptomics. Lines are coloured by menstrual phase, and each point represents pseudobulked cells assigned to a specific bin per donor. **d**, Log-normalised spatial expression of *SFRP5* on a *Visium HD* spatial transcriptomics section from proliferative phase. **e,** Log-normalised expression of *SFRP5* in *Xenium* 5K+100-plex spatial transcriptomics sections from menstrual days 2 and 5. **f**, Post-*Xenium* H&E image of a transverse section of a pre-natal uterus at 20 post-conception weeks (PCW) (top), and corresponding log-normalised spatial expression of *SFRP5* (bottom). **g,** Spatial localisation of a representative lymphoid aggregate in a *Xenium* 480-plex spatial transcriptomics section from proliferative phase (boxed region) and corresponding magnified views of *MS4A1*, *CD8A*, *CXCL13*, and *LTF* transcript localization. Visualisation generated by 10x Genomics Xenium Explorer 4.1.1. **h,** UMAP embedding of scRNA-seq cells from 11 post-menopausal donors from an independent dataset (INCLIVA) coloured by cell type annotation. **i,** Dot plot showing the expression of epithelial markers across epithelial cell subtypes identified in the post-menopausal dataset in **h**. Dot size indicates the fraction of expressing cells, and colour intensity reflects min-max normalised expression within each group. **j,** Bar plot showing the proportions of each epithelial cell subtype identified in the post-menopausal dataset in **h**. **k,** Dot plot showing the expression of the basalis stromal fibroblast marker *SFRP5* across stromal fibroblast cell subtypes identified in the post-menopausal dataset in **h**. Dot size indicates the fraction of expressing cells, and colour intensity reflects min-max normalised expression within each group. **l,** Schematic summary of proposed basalis epithelium, basalis stromal fibroblast and lymphoid aggregate markers across the menstrual cycle and in postmenopausal endometrium. Genes shown in grey in the menstrual phase and post menopausal endometrium indicate insufficient data to confidently determine their expression patterns. Pro: proliferative; Sec: secretory; Men: menstrual; ExHor: exogenous hormones.

The lower abundance of basalis glands, together with their subtle transcriptional differences relative to functionalis glands, prevented reliable separation by unsupervised clustering of *Xenium* and *Visium HD* data alone. To overcome these limitations and investigate epithelial programmes specific to basalis glands, we therefore integrated multiple orthogonal strategies. First, spatial gradient modelling across the basalis-to-luminal axis in *Xenium* 480 allowed us to identify *WIF1* and *CYP24A1* as novel markers preferentially enriched within the epithelium of deep basalis glands (**Fig. 5d**; **Extended Data Fig. 5a-d**). *WIF1*, a secreted inhibitor of Wnt signalling, showed strongest expression in basalis glands during menstruation and in exogenous hormone-treated samples, although spatial enrichment within the basalis epithelium remained detectable and specific across all menstrual phases (**Extended Data Fig. 5a-d**). Similarly, *CYP24A1* remained enriched within basalis glands throughout the menstrual cycle, but during menstruation and in exogenous hormone-treated samples its expression expanded throughout the glandular epithelium and was also detected in shed epithelial cells recovered from menstrual fluid, consistent with previous reports that *CYP24A1* is repressed by progesterone^76^ (**Extended Data Fig. 5a-d**).

Because the Xenium 480-plex panel provided limited coverage of epithelial genes, we next leveraged our *Visium HD* data of proliferative-phase sections, in which basalis glands were manually annotated prior to differential expression analysis against the surrounding tissue, to identify additional basalis-associated markers (Methods). This approach confirmed the specificity of *WIF1* and *CYP24A1*, and further identified enrichment of *SLC6A2*, *MUC6*, and *AZGP1* within basalis glands (**Fig. 5e**; **Extended Data Fig. 5a**). *SLC6A2*, which encodes the norepinephrine transporter responsible for catecholamine reuptake, was also present in the *Xenium* 5k+100 panel and showed consistent basalis-associated expression across all profiled samples (two menstrual and one exogenous hormone-treated), supporting stability of this programme across regenerative states (**Fig. 5f**, **Extended Data Fig. 5d**).

Integration of these spatially defined markers with the single-cell atlas resolved a discrete epithelial population co-expressing *WIF1*, *CYP24A1, SLC6A2*, *AZGP1*, and *MUC6* that was largely quiescent based on low expression of cell-cycle-associated genes (**Fig. 5g**, **Extended Data Fig. 5e**). Cells within this population were contributed by donors from whole-uterus samples spanning all phases of the menstrual cycle, consistent with persistence of a specialised basalis epithelial compartment across cyclical endometrial remodelling (**Extended Data Fig. 5f**).

### A Wnt-inhibitory stromal and immune niche surrounds quiescent basalis epithelium

Alongside this newly characterised *WIF1+* glandular basalis epithelial population, mapping cell type annotations onto our spatial atlas revealed basalis stromal fibroblasts and B cells which were predominantly localised to the basalis compartment across menstrual phases (**Figure 6a-b**).

The stromal fibroblast population, localised to the deepest endometrial layer surrounding basalis epithelial glands during the menstrual and proliferative phases, was characterised by specific expression of *SFRP5* (**Fig. 6c-e**; **Extended Data Fig. 6a-c**). *SFRP5* encodes a secreted antagonist of canonical and non-canonical Wnt signalling, raising the possibility that basalis stroma contributes to local suppression of Wnt activity during periods of extensive tissue remodelling, including menstrual shedding and post-menstrual regeneration. The association of *SFRP5* with progenitor-supportive mesenchymal environments was further supported by its specific expression in mesenchyme adjacent to the fetal uterine epithelium (**Fig. 6f**), and by its recent identification as a marker of lateral plate mesoderm progenitors in the human embryo^77^. Together, these findings suggest that the adult basalis retains a developmentally conserved stromal niche that may function to preserve progenitor identity during cyclical regeneration.

The basalis additionally harboured organised lymphoid aggregates enriched for *MS4A1*⁺ B cells and T cells detectable throughout the cycle, consistent with prior descriptions of such structures in the basal endometrium^78,79^ **(Fig. 6g, Extended data Fig. 6d**). Spatial resolution of these aggregates revealed colocalisation with *CXCL13*, a marker of tertiary lymphoid structures previously reported to drive B cell recruitment^80,81^, and *LTF,* which modulates B cell expansion, IgA production and T cell activity in Peyer’s patches in murine and bovine intestine^82,83^ (**Fig. 6g**). The consistent presence of these organised immune structures within the basalis, alongside the quiescent progenitor-like epithelial populations, indicates that the basalis niche may integrate both regenerative and immune surveillance functions.

The well-documented histological resemblance between atrophic postmenopausal endometrium and the basalis^38^ suggests that true basalis components should be retained in postmenopausal tissue, providing an independent context in which to validate our findings. We therefore generated single-cell transcriptomic profiles from an additional 11 postmenopausal whole-uterus samples (**Fig. 6h, Extended Data Fig. 6e**). The basalis epithelial programme was strongly enriched in postmenopausal endometrium: approximately one quarter of epithelial cells retained the *WIF1/CYP24A1/SLC6A2/AZGP1/MUC6* signature, a proportion substantially exceeding that observed in reproductive-age tissue (**Fig. 6i-j**). A subset of postmenopausal stromal cells also retained *SFRP5* expression, indicating partial persistence of the basalis stromal niche despite the absence of cyclical hormonal input (**Fig. 6k**).

Together, these findings define the endometrial basalis as an integrated, multi-component niche comprising quiescent progenitor-like epithelial cells, Wnt-inhibitory stroma, and organised lymphoid aggregates. This architecture persists throughout adult life and into the postmenopausal years, consistent with its role in maintaining long-term regenerative capacity across the reproductive lifespan (**Fig. 6l**).

## Discussion

Faithful regeneration of epithelial tissue in adult mammals is the exception rather than the rule. Villous atrophy in coeliac disease persists in 43% of adults two to five years after gluten withdrawal^84^, and incomplete regeneration is frequently accompanied by aberrant repair, such as squamous metaplasia in chronic obstructive pulmonary disease^85^ and fibrosis in idiopathic pulmonary fibrosis^86^. The human endometrium is a notable exception, completing scarless regeneration of the entire functional layer within a single menstrual cycle.

Understanding this requires a framework capturing endometrial organisation across both space and time. While spatially organised signalling gradients are a defining feature of all regenerative tissues^87^, the endometrium has a layer of complexity absent from other systems, as its architecture is cyclically remodelled under hormonal control. By combining full-thickness spatial transcriptomics with single-cell profiling of eutopic endometrium across the cycle and shed endometrial tissue, we establish continuous basalis-to-luminal transcriptional gradients as the organising principle of endometrial renewal and resolve the molecular architecture of the basalis niche. The atlas is openly available, providing a foundation for interrogating the cellular and molecular basis of endometrial health and disease.

The most distinctive feature of endometrial regeneration is its capacity to proceed scarlessly despite acute inflammation^30^. Our data suggest this paradox is resolved through spatial compartmentalisation of otherwise incompatible programmes. Building on previous work showing extensive upregulation of genes associated with ECM-remodelling in the shed endometrial compartment^88^ro-fibrotic *LRRC15⁺* myofibroblasts are confined to the shed fraction, protecting retained tissue from fibrosis-promoting signals. Inflammatory macrophages are progressively restricted to the subluminal zone as menstruation advances, spatially confining the inflammatory signal to the active breakdown region. Opposing *HSD11B1/HSD11B2* gradients partition local glucocorticoid availability across the tissue depth, generating a cortisol gradient that regionally limits inflammation^31^. Together, these mechanisms suggest the endometrium actively compartmentalises inflammatory and regenerative programmes in space to prevent fibrotic escalation. The presence of survival-programmed cells in menstrual fluid reinforces its potential as a dynamic and underutilised readout of endometrial state. Furthermore, the programmes activated in shed cells, pro-survival *BNIP5* in epithelium and pro-fibrotic *LRRC15* in stroma, may have implications beyond normal physiology: in the context of retrograde menstruation, these programmes may facilitate the ectopic survival and fibrotic remodelling that characterise endometriotic lesion establishment.

Another factor contributing to scarless repair in the endometrium is rapid re-epithelialization^64^. The expression of luminal markers *WNT7A* and *KRT17* in subluminal glands during menstrual repair suggests that glandular cells can transiently acquire luminal identity, pointing to a plastic boundary between glandular and luminal epithelium. Luminal progenitors have been identified in the endometrium^22^, with subsequent functional experiments demonstrating that WNT7A⁺ cells are required for long-term organoid survival^89^ and mouse lineage tracing confirming that luminal and glandular epithelia are maintained by distinct progenitor populations across homeostasis, menstruation, and postpartum repair^70^. This epithelial plasticity may explain why endometrial organoids can be derived from biopsies without sampling the basalis^90,91^, and raises the question of whether progenitor contributions shift across human physiological contexts, including parturition, where the urgency of re-epithelialisation differs substantially.

Within this regenerative framework, our gradient-based analysis reveals the spatiotemporal organisation of endometrial regenerative signalling. Both *IHH* and *TRH* have been proposed as markers of basalis epithelial progenitors based on sorted cell populations and immunohistochemistry^72^. Resolving this at finer spatial and temporal granularity, we show that *IHH* and *TRH* re-expand from the basalis through the glandular epithelium between menstrual days 2 and 5, pointing to the activation of an endometrial regeneration programme before resolution of bleeding. *IHH*, *PTCH1*, and *GLI1* then maintain concordant basalis-to-lumen gradients across the epithelium and stroma throughout the proliferative phase, before progesterone-mediated attenuation in the secretory phase. This demonstrates how spatiotemporal axis-aware modelling uncovers regulatory architecture which dissociated single-cell transcriptomics approaches obscure.

Leveraging our spatial atlas, we resolve a previously unrecognised basalis epithelial programme co-expressing *WIF1*, *SLC6A2*, *AZGP1*, *MUC6*, and *CYP24A1* that is largely quiescent and persists across all cycle phases. Its marked enrichment in postmenopausal endometrium, where cyclical remodelling has ceased yet regenerative capacity is retained, supports the interpretation of a long-lived reservoir maintained independently of hormonal cycling. Prior models built on individual markers such as SSEA-1^8^, N-cadherin, and AXIN2^9^ are refined by our spatial data, which show these markers extend beyond the basalis into the broader glandular epithelium, indicating regional enrichment rather than a sharp compartmental boundary. Quiescent basalis epithelial cells co-localise with a spatially restricted population of *SFRP5⁺* fibroblasts, together constituting a candidate stem cell niche distinct from the perivascular SUSD2⁺/CD146⁺ mesenchymal populations distributed throughout the functionalis^92,93^, which represent a broader mesenchymal precursor compartment rather than a localised niche-supporting population.

A defining feature of this basalis niche is local WNT regulation. Estradiol enhances WNT/β-catenin activity during the proliferative phase^94^, creating a mitogenic environment from which the basalis must be shielded to preserve progenitor quiescence. Co-expression of *WIF1* in basalis epithelial cells and *SFRP5* in surrounding stroma suggests complementary cell-autonomous and paracrine WNT inhibition. This niche architecture is shared with progenitor compartments in other glandular epithelia, where WNT inhibition prevents premature activation^95^ and *MUC6* marks basally-enriched progenitor-associated populations^96^. The presence of *SFRP5*+ stromal fibroblasts surrounding the fetal uterine epithelium further indicates that this WNT-inhibitory niche is already established during prenatal life and maintained throughout adulthood and into menopause. Moreover, *WIF1* silencing in endometrial cancer through promoter hypermethylation, resulting in aberrant WNT/β-catenin activation^97^, suggests that dissolution of this checkpoint may be an early event in malignant transformation. Definitive proof of progenitor identity of *WIF1*+ basalis epithelial cells will require functional validation, such as prospective isolation and clonogenic assays.

The spatial co-localisation of quiescent progenitor-like epithelium and organised lymphoid aggregates in the basalis is further reminiscent of gut-associated lymphoid tissue^98^. Co-expression of *CXCL13* and *LTF* within these aggregates may support immune protection of the basalis progenitors through B cell recruitment and immunoglobulin production. That endometrial aggregates reside in the basalis rather than subepithelial domes likely reflects the endometrium’s distinct immune demands compared to other mucosal tissues: interfacing with the uterine microbiome^99^ while tolerating sperm and the semi-allogeneic implanting embryo.

By mapping the spatial and molecular organisation of the regenerating endometrium, this work opens direct avenues for investigating endometrial dysfunction. HMB affects one in three women yet its endometrial origins remain incompletely understood^1,100^. Notably, *SFRP5*⁺ basalis stromal fibroblasts are significantly enriched for genetic risk variants associated with HMB in a parallel study from our group (Cohen et al. 2026, *biorxiv*), underlining the translational relevance of characterising the basalis niche. The *HSD11B1/HSD11B2* glucocorticoid axis is independently implicated in HMB by evidence that *HSD11B2* is upregulated in affected endometrium^101^, and that dexamethasone, a synthetic glucocorticoid resistant to *HSD11B2* inactivation, reduces menstrual blood loss in a randomised control trial^102^. Spatial transcriptomics could evaluate whether the glucocorticoid-partitioning H*SD11B1/HSD11B2* gradients, prostaglandin synthesis programme, and fibrinolysis inhibitor *SERPINB2* identified here are disrupted in disease. Asherman’s syndrome represents explicit failure of scarless repair following iatrogenic injury, and full-thickness spatial profiling of affected uterus would directly test whether the anti-fibrotic mechanisms described here are disrupted in the pro-fibrotic environment of intrauterine adhesions. More broadly, the endometrium offers a unique window into scarless repair that can be compared systematically with the steady-state gradients of the intestinal crypt-villus axis, wound healing in skin, and the failure of these principles in fibrotic disease.

## Methods

### Patient samples

Superficial endometrial biopsies and full-thickness samples have been collected through the “Sanger Human Cell Atlasing Project” study (Yorkshire & The Humber–Leeds East Research Ethics Committee (REC): 19/YH/0441). Menstrual fluid samples were obtained through the “Reproduction-related tissues on a cellular level” study (London – Stanmore REC: 25/LO/0357). Written informed consent was obtained from study participants before tissue samples and phenotypic data were collected. Full-thickness uterine samples were obtained from deceased transplant organ donors through the “HCA: Human Cell Atlas” study (East of England—Cambridge South REC: 15/EE/0152) and informed consent from the donor families. The uterus was removed within 1 h of circulatory arrest.

### Assignment of menstrual stage

Menstrual cycle phase was assigned using patient-reported cycle day, serum hormone measurements, and ovulation confirmation. Day 1 of the menstrual phase was defined as the first day of menstrual bleeding reported by the patient. Where discrepancies arose between these assessments, endometrial histology was additionally evaluated and the menstrual phase classified according to the Noyes criteria^103^.

### Tissue processing

Endometrial biopsy samples cryopreserved in CryoStor CS10 (STEMCELL Technologies; cat. no. 210502) were thawed at 37 °C and immediately transferred to ice-cold RPMI 1640 medium (Gibco, cat. no. 21875-034). Tissue was centrifuged (500 × g, 5 min, 4 °C) and resuspended in RPMI 1640 containing collagenase V (1 mg/mL; Sigma, cat. no. C9263), fetal bovine serum (FBS, 10% (v/v); Gibco, cat. no. A52567-01), and DNase I (0.1 mg/mL; Sigma, cat. no. 11284932001).

Samples were incubated at 37 °C on a rotator, and dissociation was monitored every 15 min for up to 1 h. When complete dissociation was achieved (i.e., no visible tissue fragments), suspensions were centrifuged, washed in DPBS without Ca²⁺ and Mg²⁺ (Gibco, cat. No. 14190144), and centrifuged at 800 × g for 2 min. Pellets were resuspended in 2 ml of red blood cell lysis buffer (ThermoFisher scientific, cat. no. 00-4333-57) and incubated for 5 min, followed by washing in DPBS and resuspension in Stromal Fibroblast Medium (DMEM, high glucose (Gibco, cat. No. 41965039), supplemented with 10% FBS (Gibco, cat. no. 10270-106) and 100 µg/ml Primocin (InvivoGen, France, cat. no. ant-pm-1)) and maintained on ice for downstream scRNA-seq.

Alternatively, samples that retained visible tissue fragments after 1 h of digestion were centrifuged at 800 × g for 2 min and resuspended in DPBS. Suspensions were passed through a 40 µm reversible strainer, with flow-through maintained on ice. Retained tissue fragments were transferred to a separate tube and subjected to TrypLE™ Select Enzyme (1X), no phenol red (Gibco, cat. no. 125630) digestion for 15 min, followed by quenching with DMEM supplemented with 10% FBS. Both fractions were centrifuged at 800 × g for 2 min and subsequently recombined in 2 ml of red blood cell lysis buffer for 5 min. Cells were then washed in DPBS and resuspended in Stromal Fibroblast Medium and maintained on ice for downstream scRNA-seq.

Menstrual fluid samples were collected from individuals without hormonal contraceptive use, using a home collection kit on days 1, 2 or 3 of the menstrual phase. Menstrual fluid was collected using a menstrual cup, transferred to a sterile 50 mL Falcon tube (Fisher Scientific, cat. no. 10203001), and dropped off at designated points on the study site within 12 h of collection. Samples were washed twice with ice-cold PBS (Gibco, cat. no. 10010015) to a final volume of 50 mL. The pellet was resuspended in 20 mL ice-cold PBS and passed through a 70 μm cell strainer to isolate tissue fragments. Tissue retained on the filter was backwashed into a dish and, where necessary, minced to ∼1 mm³ using scalpels. Tissue was enzymatically digested in RPMI 1640 medium containing collagenase V (1 mg/mL), FBS (10% (v/v)), and DNase I (0.1 mg/mL) for 20-40 min at 37 °C with rotation (10 rpm). The resulting cell suspension was further dissociated with TrypLE Express (Gibco; cat. no. 10718463) for 5-10 min at 37 °C with rotation (10 rpm), followed by quenching with PBS supplemented with FBS (10% (v/v)). Cells were incubated in red blood cell lysis buffer for 5-10 min at room temperature, filtered through a 40 μm cell strainer and collected by centrifugation (500 × g, 5 min, 4 °C). The final cell pellet was resuspended in PBS containing bovine serum albumin (BSA, 20% (v/v); Miltenyi, cat. no. 130-091-376) and EDTA (2 mM; Invitrogen, cat. no. AM9260G) prior to cell counting. Single cells were fixed and stored long-term at −80 °C until processing for single-cell RNA sequencing according to the manufacturer’s instructions (10x Genomics, GEM-X Flex Sample Preparation Kit v2, cat. no. 1000781).

Whole-uterus samples used for scRNA-seq and imaging analyses were stored in HypoThermosol FRS (STEMCELL Technologies, cat no. 101102) at 4 °C until processing. For spatial transcriptomics and imaging analyses, samples were further dissected, embedded in Optimal Cutting Temperature compound (Thermo Fisher Scientific, cat. no. 23730571), and rapidly frozen in dry ice-isopentane slurry^104^ or directly in liquid nitrogen.

### Library preparation and sequencing

#### 10x Genomics - Chromium and FLEX

Single-cell libraries on viable cells were prepared using Chromium Next GEM Single Cell 3’ v3.1 (10x Genomics; cat no. 1000268), Chromium GEM-X Single Cell 3’ v4 (10x Genomics; cat no. 1000691) or Chromium GEM-X Single Cell 5’ v3 (10x Genomics; cat no. 1000699) according to the manufacturer’s instructions, aiming to obtain 5,000-10,000 cells per reaction. All libraries were sequenced on the NovaSeq 6000 platform, targeting a minimum of 50,000 reads per cell, using the sequencing format: Read 1: 28 cycles; i7 index: 8 cycles; i5 index: 0 cycles; Read 2: 98 cycles.

Single-cell libraries on fixed cells were prepared using the GEM-X Flex Gene Expression Human kit (10x Genomics; cat no. 1000793) according to the manufacturer’s instructions, aiming to obtain 20,000 cells per reaction. All libraries were sequenced on the NovaSeq X platform with 1.5b or 10b flow cells, targeting a minimum of 50,000 reads per cell, using the sequencing format: Read 1: 28 cycles; i7 index: 10 cycles; i5 index: 10 cycles; Read 2: 90 cycles.

#### 10x Genomics - Xenium

Spatial transcriptomic profiling was performed on fresh frozen tissue sections using the 10x Genomics Xenium In Situ platform. Tissue blocks were embedded in Optimal Cutting Temperature (OCT) compound immediately following collection and frozen as above. Cryosections of 10 µm thickness were placed onto Xenium slides, fixed, and permeabilised following the manufacturer’s Fresh Frozen Tissue Preparation Handbook (CG000579, Rev D) and Fixation & Permeabilization Demonstrated Protocol (CG000581).

Two panel configurations were applied across specimens. The first used a fully bespoke 480-gene standalone custom panel (design ID JW4ZDU). The second used the Xenium Prime 5K pre-designed panel supplemented with a 100-gene custom add-on (design ID JEJN79 and PYDA6C), both configured following the Xenium Panel Designer Advanced Workflows Guide. The fully custom probe panel was designed to target 480 genes that capture the diversity of cell states in the human endometrium across phases of the menstrual cycle (Supplementary Table 2). Genes were selected from our previously published human endometrial cell atlas^21^and were complemented by markers previously implicated in endometrial biology and endometrial conditions from the literature.

Probe hybridisation, washing, ligation, amplification, and cell segmentation staining were performed following the Xenium In Situ Gene Expression with Cell Segmentation Staining User Guide (CG000749) for the 480-gene custom panel, and the Xenium Prime In Situ Gene Expression with added Cell Segmentation Staining User Guide (CG000760) for the 5K + 100-gene add-on panel. Imaging and transcript decoding were carried out on the Xenium Analyzer (CG000584), with nuclear detection, multimodal cell segmentation, and transcript deduplication performed by Xenium Onboard Analysis (XOA v4.0). Post-instrument H&E staining followed (CG000613). Data were re-analysed using Xenium Ranger v4.0.1.0.

### 10x Genomics Visium HD

Whole-transcriptome spatial gene expression profiling was performed on fresh frozen tissue sections using the 10x Genomics Visium HD platform with CytAssist. Sections of 10 µm thickness were cut with a cryostat, and tissue block preparation, sectioning, H&E staining, and imaging followed the Visium HD Fresh Frozen Tissue Preparation Handbook (CG000763). Probe hybridisation, probe ligation, slide preparation, probe release, extension, and library construction followed the Visium HD Spatial Gene Expression Reagent Kits User Guide (CG000685). Probe transfer from tissue slides to Visium HD capture slides was performed using the Visium CytAssist instrument (firmware v2.0.0 or higher), following the Tissue Slide Alignment Quick Reference Cards (CG000548). Libraries were sequenced on NovaSeqX.

Space Ranger v4.01 was used to map reads to the reference, detect tissue, align data to the microscope and CytAssist images, and output feature-barcode matrices for further analysis.

### Single-cell RNA sequencing quality control

Raw sequencing data were processed using Cell Ranger (10x Genomics v9.1, including introns) to generate cell by gene count matrices aligned to the GRCh38 reference genome. Ambient RNA contamination was removed from each count matrix using CellBender v0.3.2^105^, run per sample with default parameters.

Per-cell quality control metrics were computed independently for each library, including the number of detected genes, total UMI counts, and mitochondrial read fraction. Doublets were identified using Scrublet. Additional per-cell scores were computed to capture biological and technical variation, including a senescence score and a dissociation-induced stress score^106^. Cells were filtered based on the joint distribution of these metrics; outlier libraries and clusters with elevated mitochondrial fractions, low complexity, or high doublet scores were removed. Cell cycle phase (G1, S, G2/M) was assigned using the Scanpy score genes of previously defined cell cycle genes^107^.

### scRNA-seq integration and annotation

Highly variable genes (HVGs, n = 3,000) were selected using the Seurat v3 method, restricted to genes expressed in at least 5 cells. Dimensionality reduction and batch integration were performed using scVI^108^Gene expression counts were modelled using a negative binomial likelihood with a gene-specific inverse dispersion parameter. The dataset of origin was specified as the batch covariate, and donor identity and cell cycle phase were included as additional categorical covariates in the design matrix. The encoder and decoder comprised one hidden layer, and the dimensionality of the latent space was set to 60. Models were trained for a maximum of 200 epochs with early stopping. Leiden clustering on the resulting latent space was used to assign broad lineage labels (mesenchymal, epithelial, endothelial, immune, and neural) based on a curated panel of canonical marker genes.

Cells from each lineage were subsequently subset and analysed independently. Within each lineage, HVGs were selected using the intersection of genes across datasets, and used to train scVI models with the architecture described above, with a gene- and batch-specific dispersion parameter. The resulting latent representation was used for downstream analyses.

Cluster marker genes were identified using term frequency–inverse document frequency (TF-IDF) as implemented in SoupX^109^ (using 100 cells downsampled per cluster, top 50 markers retained at FDR < 0.01). Cluster identities were assigned by integrating TF-IDF markers, canonical marker gene expression, and metadata distributions (tissue compartment, donor, menstrual cycle stage). Low-quality clusters identified during annotation were excluded from the final object.

For downstream gene expression analysis, counts were normalised to 10,000 counts per cell and log1p-transformed.

### Preprocessing of Visium HD data and manual annotation of basalis glands

Visium HD datasets were preprocessed using bin2cell, which aggregates neighbouring 2 µm bins assigned to the same segmented object to generate cell-level profiles while preserving spatial coordinates and high-resolution morphological information for visualisation^110^. Basalis epithelial glands were then manually annotated using TissueTag2, enabling precise delineation of glandular structures within the deepest endometrial layer^41^. To identify basalis-associated epithelial programmes, we performed differential expression analysis between manually annotated glands and all remaining tissue using a TF-IDF-based framework, thereby enriching for genes specifically associated with basalis glandular epithelium relative to the surrounding endometrial microenvironment.

### Post-Xenium H&E registration and anatomical landmark annotation

Following completion of each Xenium run, post-run haematoxylin and eosin (H&E)-stained sections were imaged at 40× magnification. H&E images were converted to OME-TIFF format and registered to the DAPI-stained Xenium image using the 10x Genomics Xenium Explorer software.

Anatomical compartment boundaries were annotated on each registered H&E image in a two-stage workflow. First, H&E images were tiled at native resolution (0.22 µm/pixel), and morphological embeddings were extracted for each tile using UNI2, a computational pathology foundation model based on a ViT-H architecture pre-trained on >100,000 histological images^40^. Tiles were clustered in embedding space, and cluster assignments were projected back onto the H&E image to generate an unsupervised pre-annotation map partitioning tissue into morphologically coherent regions.

Second, the UNI2-derived cluster maps were imported into TissueTag2^41^, an interactive Jupyter-based image annotation tool, for manual review and refinement of compartment boundaries. Four anatomical compartment labels were assigned: (i) myometrium; (ii) basalis, the deeper endometrial layer characterised by horizontally oriented glands; (iii) functionalis, the superficial endometrial layer characterised by vertically oriented glands; and (iv) luminal epithelium, the continuous single-cell layer lining the uterine cavity.

### Construction of the normalised basalis-to-luminal axis

A universal positional axis spanning the full endometrial thickness was constructed to enable comparison of cell positions across sections with variable compartment thickness arising from cycle phase, pathology, or incomplete tissue preservation. The axis was defined on the interval [0,1], where 0 denotes the myometrial-endometrial junction and 1 denotes the luminal surface. Basalis occupied the interval [0,p], and functionalis occupied [p,1].

Pixel-based compartment annotations were converted to a hexagonal grid of spots at 10 µm spacing using TissueTag2^41^. Euclidean distances from each grid spot to adjacent compartment boundaries were then computed. Within-compartment fractional position was calculated from these boundary distances. For basalis spots, fractional position was defined as the distance from the myometrial boundary divided by the sum of distances from the myometrial and basalis–functionalis boundaries. For functionalis spots, the analogous ratio was computed using distances from the basalis-functionalis and luminal boundaries. This local normalisation defines position independently of absolute compartment thickness at each tissue location.

Within-compartment fractions were mapped onto the universal axis using a cohort-derived compartment proportion, p, estimated as the median ratio of basalis thickness to total endometrial thickness across complete sections containing intact basalis, functionalis, and luminal epithelium. Per-sample compartment thickness was estimated as the 95th percentile of local thickness measurements across grid spots to reduce sensitivity to edge artefacts.

Sections lacking luminal epithelium (<50 annotated grid spots) were classified as truncated. In these sections, expected full functionalis thickness was imputed from the observed basalis thickness and the cohort-derived median basalis:functionalis thickness ratio. Positional coordinates for functionalis cells were then estimated from distance to the basalis boundary alone.

Each annotated compartment (basalis, functionalis, lumen) was subdivided into spatial bins along its within-compartment axis using equal-width intervals: the basalis into two bins (basalis_1, basalis_2) and the functionalis into three (functionalis_1, functionalis_2, functionalis_3). This design choice aims to approximately reflect the 1:2 proportional depth of the basalis and functionalis layers in the endometrium^111^. The luminal epithelium was retained as a single bin (lumen_1). Bin boundaries were defined separately within each annotated compartment rather than across the full axis, so they represent equal divisions of each compartment’s local axis range rather than absolute positions. The resulting six bins, ordered basalis_1 through lumen_1, constitute computational intervals along a continuous spatial gradient and do not correspond to discrete anatomical boundaries.

### Spatial axis mapping onto scRNA-seq reference

To assign spatial positional information to cells in the scRNA-seq reference atlas, we transferred the myometrial-luminal axis defined above by computationally pairing cells from the two modalities using either ISS-patcher^42^ or TACCO^43^ (**Supplementary Note**). Both methods were run with default parameters. All transfer was performed separately for epithelial and stromal compartments, and restricted to stage-matched cells, whereby single-cell profiles were paired within the same menstrual phase. Menstrual fluid samples were excluded from this analysis.

To identify genes with spatially organised expression along the basalis-to-lumen axis, single-cell profiles were aggregated into pseudobulk samples per donor per spatial compartment. Pseudobulk observations with fewer than 20 cells were excluded, and donors contributing fewer than two valid bins after filtering were removed. Due to a low number of epithelial cells from the menstrual stage, analysis was restricted to proliferative and secretory phase donors for the epithelial lineage. Pseudobulk expression was quantified as log₂(CPM + 1), where library sizes were computed per pseudobulk sample. Prior to modelling, genes were retained if they exceeded log₂(CPM + 1) ≥ 1 in at least 10 pseudobulk samples. Mitochondrial genes and ribosomal protein genes were removed. The bottom 25% of genes by cross-sample variance were also excluded from the input modelling gene set.

For each retained gene, two linear mixed models with Gaussian errors were fitted by maximum likelihood using statsmodels^112^: a reduced model including axis position and menstrual phase as fixed effects, and a full model additionally including their interaction. Axis position was encoded as a continuous integer (basalis_1 = 1, …, lumen_1 = 6), and a per-donor random intercept was included in both models to account for inter-individual variation. Axis-associated genes (Test A) were identified via Wald test on the axis coefficient from the reduced model, with beta representing the per-bin change in log₂-CPM from basalis to lumen. Axis-phase interaction effects (Test B) were assessed by likelihood ratio test comparing the full and reduced models. P-values from each test were corrected independently using the Benjamini-Hochberg procedure at FDR < 0.05

## Data and code availability

We provide user-friendly access to our annotated scRNA-seq resource via cellxgene at https://www.reproductivecellatlas.org/HCAreproductive/v1/uterus/. All the raw and processed sequencing data generated in this study are currently being deposited to EGA and BioImageArchive (10x Xenium). The code used to perform the analyses presented in the manuscript can be found at https://github.com/ventolab/Human-Uterine-Spatial-Cell-Atlas.

## Acknowledgements

We acknowledge Jackie Maybin, program director, and Alessandro Chioino, program coordinator from the Wellcome Leap Missed Vital Sign program for their support. This work was created in collaboration with the Faculty of Medicine, Masaryk University, within the CZECRIN project (LM2023049), supported by the Ministry of Education, Youth and Sports from the state budget of Czech Republic. We thank the transplant organ donors and their families for the samples donated through the Cambridge Biorepository for Translational Medicine. We also thank Júlia Fernández-Lechiguero and the Departments of Obstetrics and Gynecology and Pathology of the General University Hospital of Valencia for patient recruitment and clinical sample collection. We thank Hongorzul Davaapil, Anna Wallis, Kamila Marciniak, and Georgia Leisegang in the Sanger Cellular Services Core Facility and the Sanger Core Sequencing pipeline for sample processing and sequencing library preparation; The Scientific Operations Spatial team for spatial sample profiling; A. Predeus for assistance with reprocessing public scRNA-seq datasets; T. Porter, E. Tuck, and the Cellular Genetics wet lab team for experimental support; A. García of Bio-Graphics for scientific illustrations; and A. Maartens for proofreading. This publication is part of the Human Cell Atlas – www.humancellatlas.org/publications/.

## Author information

V.L and R.V-T conceived and designed the experiments and analyses. V.L and M.M analysed the data with contributions from C.E.C, A.P, C.E.K, R.S, M.P and N.Y. R.C, M.D, and C.S-S performed sample processing for scRNA-seq with contributions from C.C under the supervision of I.K. C.I.M performed sample processing for spatial transcriptomics. J.H, K.C, M.V, A.H, D.R, M.T, J.M.A, S.T, C.A, K.T.A., K.S-P, T.G, I.M, A.M. contributed to endometrial sample acquisition supervised by F.V, V.W, V.P-G, J-G-E, C.S, S.A.T. V.L, M.M, L.G-A, and R.V-T interpreted the data. V.L, M.M, and R.V-T wrote the manuscript with contributions from L.G-A. R.V-T supervised the work. All authors read and approved the manuscript.

## References

1. Jain, V., Chodankar, R. R., Maybin, J. A. & Critchley, H. O. D. Uterine bleeding: how understanding endometrial physiology underpins menstrual health. Nat Rev Endocrinol 18, 290–308 (2022).

2. Zondervan, K. T. et al. Endometriosis. Nat Rev Dis Primers 4, 9 (2018).

3. Santamaria, X. et al. Decoding the endometrial niche of Asherman’s Syndrome at single-cell resolution. Nat Commun 14, 5890 (2023).

4. Santamaria, X., Isaacson, K. & Simón, C. Asherman’s Syndrome: it may not be all our fault. Hum. Reprod. 33, 1374–1380 (2018).

5. Chan, R. W. S., Schwab, K. E. & Gargett, C. E. Clonogenicity of human endometrial epithelial and stromal cells. Biol. Reprod. 70, 1738–1750 (2004).

6. Gargett, C. E., Schwab, K. E., Zillwood, R. M., Nguyen, H. P. T. & Wu, D. Isolation and culture of epithelial progenitors and mesenchymal stem cells from human endometrium. Biol. Reprod. 80, 1136–1145 (2009).

7. Cervelló, I. et al. Human endometrial side population cells exhibit genotypic, phenotypic and functional features of somatic stem cells. PLoS One 5, e10964 (2010).

8. Valentijn, A. J. et al. SSEA-1 isolates human endometrial basal glandular epithelial cells: phenotypic and functional characterization and implications in the pathogenesis of endometriosis. Hum Reprod 28, 2695–2708 (2013).

9. Nguyen, H. P. T. et al. N-cadherin identifies human endometrial epithelial progenitor cells by in vitro stem cell assays. Hum. Reprod. 32, 2254–2268 (2017).

10. Jin, S. Bipotent stem cells support the cyclical regeneration of endometrial epithelium of the murine uterus. Proc. Natl. Acad. Sci. U. S. A. 116, 6848–6857 (2019).

11. Syed, S. M. et al. Endometrial Axin2+ cells drive epithelial homeostasis, regeneration, and cancer following oncogenic transformation. Cell Stem Cell 26, 64–80.e13 (2020).

12. Zhang, Y. et al. A spatial transcriptomics dataset of the endometrium from repeated implantation failure patients. Sci Data 12, 1711 (2025).

13. Tempest, N., Soul, J., Hill, C. J., Caamaño Gutierrez, E. & Hapangama, D. K. Cell type and region-specific transcriptional changes in the endometrium of women with RIF identify potential treatment targets. Proc Natl Acad Sci U S A 122, e2421254122 (2025).

14. Liu, Y. et al. Decrypting spatiotemporal code of human endometrial receptivity. Nat Commun 17, 801 (2025).

15. Burns, G. W. et al. Single-cell mapping of human endometrium and decidua reveals epithelial and stromal contributions to fertility. JCI Insight 11, (2026).

16. Liu, S. et al. Single-cell and spatial transcriptomic profiling revealed niche interactions sustaining growth of endometriotic lesions. Cell Genom 5, 100737 (2025).

17. Wang, W. et al. Single-cell transcriptomic atlas of the human endometrium during the menstrual cycle. Nat. Med. 26, 1644–1653 (2020).

18. Tan, Y. et al. Single-cell analysis of endometriosis reveals a coordinated transcriptional programme driving immunotolerance and angiogenesis across eutopic and ectopic tissues. Nat Cell Biol 24, 1306–1318 (2022).

19. Huang, X. et al. Single-cell transcriptome analysis reveals endometrial immune microenvironment in minimal/mild endometriosis. Clin Exp Immunol 212, 285–295 (2023).

20. Ulrich, N. D. et al. Cellular heterogeneity and dynamics of the human uterus in healthy premenopausal women. Proc Natl Acad Sci U S A 121, e2404775121 (2024).

21. Marečková, M. et al. An integrated single-cell reference atlas of the human endometrium. Nat. Genet. 56, 1925–1937 (2024).

22. Garcia-Alonso, L. et al. Mapping the temporal and spatial dynamics of the human endometrium in vivo and in vitro. Nat. Genet. 53, 1698–1711 (2021).

23. Zhang, Y. et al. A spatial transcriptomics dataset of the endometrium from repeated implantation failure patients. Sci Data 12, 1711 (2025).

24. Tempest, N., Soul, J., Hill, C. J., Caamaño Gutierrez, E. & Hapangama, D. K. Cell type and region-specific transcriptional changes in the endometrium of women with RIF identify potential treatment targets. Proc Natl Acad Sci U S A 122, e2421254122 (2025).

25. Liu, Y. et al. Decrypting spatiotemporal code of human endometrial receptivity. Nat Commun 17, 801 (2025).

26. Watters, M., Martínez-Aguilar, R. & Maybin, J. A. The menstrual endometrium: From physiology to future treatments. *Front*. Reprod. Health 3, 794352 (2021).

27. Evans, J. & Salamonsen, L. A. Inflammation, leukocytes and menstruation. Rev Endocr Metab Disord 13, 277–288 (2012).

28. Garry, R., Hart, R., Karthigasu, K. A. & Burke, C. A re-appraisal of the morphological changes within the endometrium during menstruation: a hysteroscopic, histological and scanning electron microscopic study. Hum Reprod 24, 1393–1401 (2009).

29. Ludwig, H. & Spornitz, U. M. Microarchitecture of the human endometrium by scanning electron microscopy: menstrual desquamation and remodeling. Ann N Y Acad Sci 622, 28–46 (1991).

30. Critchley, H. O. D., Maybin, J. A., Armstrong, G. M. & Williams, A. R. W. Physiology of the Endometrium and Regulation of Menstruation. Physiological Reviews 100, 1149–1179 (2020).

31. Maybin, J. A. & Critchley, H. O. D. Menstrual physiology: implications for endometrial pathology and beyond. Hum. Reprod. Update 21, 748–761 (2015).

32. Maybin, J. A. et al. Hypoxia and hypoxia inducible factor-1α are required for normal endometrial repair during menstruation. Nat. Commun. 9, 295 (2018).

33. Thiruchelvam, U. et al. Cortisol regulates the paracrine action of macrophages by inducing vasoactive gene expression in endometrial cells. J. Leukoc. Biol. 99, 1165–1171 (2016).

34. Maybin, J. A., Hirani, N., Jabbour, H. N. & Critchley, H. O. D. Novel roles for hypoxia and prostaglandin E2 in the regulation of IL-8 during endometrial repair. Am. J. Pathol. 178, 1245–1256 (2011).

35. Schwalie, P. C. et al. Single-cell characterization of menstrual fluid at homeostasis and in endometriosis. eLife (2024) doi:10.7554/elife.99558.1.

36. Shih, A. J. et al. Single-cell analysis of menstrual endometrial tissues defines phenotypes associated with endometriosis. BMC Med. 20, 315 (2022).

37. Ulrich, N. D. et al. Cellular heterogeneity and dynamics of the human uterus in healthy premenopausal women. Proc. Natl. Acad. Sci. U. S. A. 121, e2404775121 (2024).

38. Ferenczy, A. & Bergeron, C. Histology of the human endometrium: from birth to senescence. Ann. N. Y. Acad. Sci. 622, 6–27 (1991).

39. Nguyen, H. P. T., Sprung, C. N. & Gargett, C. E. Differential expression of Wnt signaling molecules between pre- and postmenopausal endometrial epithelial cells suggests a population of putative epithelial stem/progenitor cells reside in the basalis layer. Endocrinology 153, 2870–2883 (2012).

40. Chen, R. J. et al. Towards a general-purpose foundation model for computational pathology. Nat Med 30, 850–862 (2024).

41. *TissueTag2: Python Package to Interactively Annotate Histological Images within a Jupyter Notebook*. (Github).

42. To, K. et al. A multi-omic atlas of human embryonic skeletal development. Nature 635, 657–667 (2024).

43. Mages, S. et al. TACCO unifies annotation transfer and decomposition of cell identities for single-cell and spatial omics. Nat Biotechnol 41, 1465–1473 (2023).

44. Gaide Chevronnay, H. P., et al. Regulation of matrix metalloproteinases activity studied in human endometrium as a paradigm of cyclic tissue breakdown and regeneration. Biochim Biophys Acta 1824, 146–156 (2012).

45. Thiruchelvam, U., Dransfield, I., Saunders, P. T. K. & Critchley, H. O. D. The importance of the macrophage within the human endometrium. J Leukoc Biol 93, 217–225 (2013).

46. Yang, H.-L. et al. Decidual stromal cells promote the differentiation of CD56 CD16 NK cells by secreting IL-24 in early pregnancy. Am J Reprod Immunol 81, e13110 (2019).

47. Tu, T. et al. Proinflammatory macrophages release CXCL5 to regulate T cell function and limit effects of αPD-1 in steatosis-driven liver cancer. JHEP Rep 7, 101385 (2025).

48. Lee, S. H., et al. M2-like, dermal macrophages are maintained via IL-4/CCL24-mediated cooperative interaction with eosinophils in cutaneous leishmaniasis. Sci Immunol 5, (2020).

49. Dollt, C. et al. The novel immunoglobulin super family receptor SLAMF9 identified in TAM of murine and human melanoma influences pro-inflammatory cytokine secretion and migration. Cell Death Dis 9, 939 (2018).

50. Vasamsetti, S. B. et al. Tissue-resident macrophage survival depends on mitochondrial function regulated by SerpinB2 in chronic inflammation. Nat Commun 17, 1493 (2026).

51. Medcalf, R. L. Plasminogen activator inhibitor type 2: still an enigmatic serpin but a model for gene regulation. Methods Enzymol 499, 105–134 (2011).

52. Law, A. Y. S. et al. Epigenetic and HIF-1 regulation of stanniocalcin-2 expression in human cancer cells. Exp Cell Res 314, 1823–1830 (2008).

53. Lv, H. et al. Mesenchymal stromal cells ameliorate acute lung injury induced by LPS mainly through stanniocalcin-2 mediating macrophage polarization. Ann. Transl. Med. 8, 334 (2020).

54. Lee, C. G. et al. Role of chitin and chitinase/chitinase-like proteins in inflammation, tissue remodeling, and injury. Annu Rev Physiol 73, 479–501 (2011).

55. Krishnamurty, A. T. et al. LRRC15 myofibroblasts dictate the stromal setpoint to suppress tumour immunity. Nature 611, 148–154 (2022).

56. Moss, A. et al. Origins, actions and dynamic expression patterns of the neuropeptide VGF in rat peripheral and central sensory neurones following peripheral nerve injury. Mol Pain 4, 62 (2008).

57. Kanehisa, K. et al. Neuronal pentraxin 2 is required for facilitating excitatory synaptic inputs onto spinal neurons involved in pruriceptive transmission in a model of chronic itch. Nat Commun 13, 2367 (2022).

58. Zimmer, A. et al. Hypoalgesia in mice with a targeted deletion of the tachykinin 1 gene. Proc Natl Acad Sci U S A 95, 2630–2635 (1998).

59. Ogami, T. et al. MAB21L4 regulates the TGF-β-induced expression of target genes in epidermal keratinocytes. J Biochem 171, 399–410 (2022).

60. Kvedar, J. C., Manabe, M., Phillips, S. B., Ross, B. S. & Baden, H. P. Characterization of sciellin, a precursor to the cornified envelope of human keratinocytes. Differentiation 49, 195–204 (1992).

61. Zheng, H. et al. SerpinB7 deficiency contributes to development of psoriasis via calcium-mediated keratinocyte differentiation dysfunction. Cell Death Dis 13, 635 (2022).

62. Vossenaar, E. R., Zendman, A. J. W., van Venrooij, W. J. & Pruijn, G. J. M. PAD, a growing family of citrullinating enzymes: genes, features and involvement in disease. Bioessays 25, 1106–1118 (2003).

63. Ruehl, S. et al. Anti-apoptotic BH3-only proteins inhibit Bak-dependent apoptosis. bioRxiv 2022.07.24.499430 (2022) doi:10.1101/2022.07.24.499430.

64. Salamonsen, L. A. Menstrual fluid factors mediate endometrial repair. *Front*. Reprod. Health 3, 779979 (2021).

65. Ko, C. J., Cho, Y. M., Ham, E., Cacioppo, J. A. & Park, C. J. Endothelin 2: a key player in ovulation and fertility. Reproduction 163, R71–R80 (2022).

66. Niethammer, P., Grabher, C., Look, A. T. & Mitchison, T. J. A tissue-scale gradient of hydrogen peroxide mediates rapid wound detection in zebrafish. Nature 459, 996–999 (2009).

67. Razzell, W., Evans, I. R., Martin, P. & Wood, W. Calcium flashes orchestrate the wound inflammatory response through DUOX activation and hydrogen peroxide release. Curr Biol 23, 424–429 (2013).

68. Nikolakopoulou, K. et al. An in vitro menstrual cycle using organoids captures epithelial cell transitions during menstruation and regeneration of the human endometrium. Cell Stem Cell 33, 747–762.e8 (2026).

69. Fukuhara, S. et al. The sphingosine-1-phosphate transporter Spns2 expressed on endothelial cells regulates lymphocyte trafficking in mice. J Clin Invest 122, 1416–1426 (2012).

70. Ang, C. J., et al. Luminal epithelium remodeling underlies endometrial regeneration during menstruation and pregnancy. *bioRxiv* (2026) doi:10.64898/2026.03.08.710375.

71. Squair, J. W. et al. Confronting false discoveries in single-cell differential expression. Nat. Commun. 12, 5692 (2021).

72. Fitzgerald, H. et al. Molecular signature of human endometrial stem/progenitor cells at the single cell level. bioRxiv 2025.06.23.660982 (2025) doi:10.1101/2025.06.23.660982.

73. Wei, Q., Levens, E. D., Stefansson, L. & Nieman, L. K. Indian Hedgehog and Its Targets in Human Endometrium: Menstrual Cycle Expression and Response to CDB-2914. The Journal of Clinical Endocrinology & Metabolism 95, 5330–5337 (2010).

74. Lv, H. et al. Deciphering the endometrial niche of human thin endometrium at single-cell resolution. Proc Natl Acad Sci U S A 119, (2022).

75. Huang, S. et al. Suppression of OGN in lung myofibroblasts attenuates pulmonary fibrosis by inhibiting integrin αv-mediated TGF-β/Smad pathway activation. Matrix Biol. 132, 87–97 (2024).

76. Bokhari, A. A. et al. Progesterone potentiates the growth inhibitory effects of calcitriol in endometrial cancer via suppression of CYP24A1. Oncotarget 7, 77576–77590 (2016).

77. Webb, S. et al. An integrated single cell and spatial omics atlas of human prenatal development. bioRxiv 2026.03.30.714220 (2026) doi:10.64898/2026.03.30.714220.

78. Shen, M. et al. The role of endometrial B cells in normal endometrium and benign female reproductive pathologies: a systematic review. Hum. Reprod. Open 2022, hoab043 (2022).

79. Yeaman, G. R. et al. Unique CD8+ T cell-rich lymphoid aggregates in human uterine endometrium. J. Leukoc. Biol. 61, 427–435 (1997).

80. Cabrita, R. et al. Tertiary lymphoid structures improve immunotherapy and survival in melanoma. Nature 577, 561–565 (2020).

81. Rodriguez, A. B. et al. Immune mechanisms orchestrate tertiary lymphoid structures in tumors via cancer-associated fibroblasts. Cell Rep. 36, 109422 (2021).

82. Miyauchi, H., Kaino, A., Shinoda, I., Fukuwatari, Y. & Hayasawa, H. Immunomodulatory effect of bovine lactoferrin pepsin hydrolysate on murine splenocytes and Peyer’s patch cells. J. Dairy Sci. 80, 2330–2339 (1997).

83. Arciniega-Martínez, I. M. et al. Modulatory effects of oral bovine lactoferrin on the IgA response at inductor and effector sites of distal small intestine from BALB/c mice. Arch. Immunol. Ther. Exp. (Warsz.) 64, 57–63 (2016).

84. Lebwohl, B., Murray, J. A., Rubio-Tapia, A., Green, P. H. R. & Ludvigsson, J. F. Predictors of persistent villous atrophy in coeliac disease: a population-based study. Aliment. Pharmacol. Ther. 39, 488–495 (2014).

85. Puchelle, E., Zahm, J.-M., Tournier, J.-M. & Coraux, C. Airway epithelial repair, regeneration, and remodeling after injury in chronic obstructive pulmonary disease. Proc. Am. Thorac. Soc. 3, 726–733 (2006).

86. Moss, B. J., Ryter, S. W. & Rosas, I. O. Pathogenic mechanisms underlying idiopathic pulmonary fibrosis. Annu. Rev. Pathol. 17, 515–546 (2022).

87. Kosinski, C. et al. Gene expression patterns of human colon tops and basal crypts and BMP antagonists as intestinal stem cell niche factors. Proc Natl Acad Sci U S A 104, 15418–15423 (2007).

88. Gaide Chevronnay, H. P., et al. Spatiotemporal coupling of focal extracellular matrix degradation and reconstruction in the menstrual human endometrium. Endocrinology 150, 5094–5105 (2009).

89. Nikolakopoulou, K. et al. An in vitro menstrual cycle using organoids captures epithelial cell transitions during menstruation and regeneration of the human endometrium. Cell Stem Cell 33, 747–762. e8 (2026).

90. Boretto, M. et al. Development of organoids from mouse and human endometrium showing endometrial epithelium physiology and long-term expandability. Development 144, 1775–1786 (2017).

91. Turco, M. Y. et al. Long-term, hormone-responsive organoid cultures of human endometrium in a chemically defined medium. Nat. Cell Biol. 19, 568–577 (2017).

92. Schwab, K. E. & Gargett, C. E. Co-expression of two perivascular cell markers isolates mesenchymal stem-like cells from human endometrium. Hum Reprod 22, 2903–2911 (2007).

93. Masuda, H., Anwar, S. S., Bühring, H.-J., Rao, J. R. & Gargett, C. E. A novel marker of human endometrial mesenchymal stem-like cells. Cell Transplant 21, 2201–2214 (2012).

94. Hou, X., Tan, Y., Li, M., Dey, S. K. & Das, S. K. Canonical Wnt signaling is critical to estrogen-mediated uterine growth. Mol. Endocrinol. 18, 3035–3049 (2004).

95. Clevers, H. & Nusse, R. Wnt/β-catenin signaling and disease. Cell 149, 1192–1205 (2012).

96. Lavery, D. L. et al. The stem cell organisation, and the proliferative and gene expression profile of Barrett’s epithelium, replicates pyloric-type gastric glands. Gut 63, 1854–1863 (2014).

97. Zhang, B. et al. WIF1 promoter hypermethylation induce endometrial carcinogenesis through the Wnt/β-catenin signaling pathway. Am. J. Reprod. Immunol. 90, e13743 (2023).

98. Bemark, M., Pitcher, M. J., Dionisi, C. & Spencer, J. Gut-associated lymphoid tissue: a microbiota-driven hub of B cell immunity. Trends Immunol. 45, 211–223 (2024).

99. Toson, B., Simon, C. & Moreno, I. The endometrial microbiome and its impact on human conception. Int. J. Mol. Sci. 23, 485 (2022).

100. Schoep, M. E., Nieboer, T. E., van der Zanden, M., Braat, D. D. M. & Nap, A. W. The impact of menstrual symptoms on everyday life: a survey among 42,879 women. Am. J. Obstet. Gynecol. 220, 569.e1–569.e7 (2019).

101. Rae, M. et al. Cortisol inactivation by 11beta-hydroxysteroid dehydrogenase-2 may enhance endometrial angiogenesis via reduced thrombospondin-1 in heavy menstruation. J. Clin. Endocrinol. Metab. 94, 1443–1450 (2009).

102. Warner, P. et al. Low dose dexamethasone as treatment for women with heavy menstrual bleeding: A response-adaptive randomised placebo-controlled dose-finding parallel group trial (DexFEM). EBioMedicine 69, 103434 (2021).

103. Noyes, R. W., Hertig, A. T. & Rock, J. Dating the endometrial biopsy. Am J Obstet Gynecol 122, 262–263 (1975).

104. Roberts, K. & Tuck, L. Embedding and freezing fresh human tissue in OCT using isopentane V.3. protocols.io https://www.protocols.io/view/embedding-and-freezing-fresh-human-tissue-in-oct-u-bp2l64o55vqe/v3 (2019).

105. Fleming, S. J. et al. Unsupervised removal of systematic background noise from droplet-based single-cell experiments using CellBender. Nat. Methods 20, 1323–1335 (2023).

106. van den Brink, S. C., et al. Single-cell sequencing reveals dissociation-induced gene expression in tissue subpopulations. Nat. Methods 14, 935–936 (2017).

107. Tirosh, I. et al. Single-cell RNA-seq supports a developmental hierarchy in human oligodendroglioma. Nature 539, 309–313 (2016).

108. Lopez, R., Regier, J., Cole, M. B., Jordan, M. I. & Yosef, N. Deep generative modeling for single-cell transcriptomics. Nat. Methods 15, 1053–1058 (2018).

109. Young, M. D. & Behjati, S. SoupX removes ambient RNA contamination from droplet-based single-cell RNA sequencing data. Gigascience 9, giaa151 (2020).

110. Polański, K. et al. Bin2cell reconstructs cells from high resolution Visium HD data. Bioinformatics 40, (2024).

111. Figueira, P. G. M., Abrão, M. S., Krikun, G. & Taylor, H. S. Stem cells in endometrium and their role in the pathogenesis of endometriosis: Stem cells, the endometrium, and endometriosis. Ann. N. Y. Acad. Sci. 1221, 10–17 (2011).

112. statsmodels 0.14.4. https://www.statsmodels.org/stable/index.html

