## Supplementary Figures for "Full-thickness spatial transcriptomics of the human uterus reveals basalis niche architecture and regeneration gradients during menstrual breakdown"

### Supplementary Note: pairing single cell and spatial transcriptomics

#### Cell type mapping onto query spatial transcriptomics samples

To assign cell type identities to segmented cells in the query spatial transcriptomics samples, we transferred cell type labels defined in the scRNA-seq reference atlas by computationally pairing cells from the two modalities, considering two possible algorithms. First, we applied ISS-patcher [1] to assign each spatial observation the cell type label of its k-nearest neighbors in the single-cell reference (here k = 30), either using the full reference or downsampling to 1,000 cells per cell type. Second, we applied TACCO[2], which uses optimal transport to quantify how much each cell type in the reference contributes to an observation in the query, here assigning contribution scores across all reference cell types to each spatial transcriptome. The cell type of highest predicted contribution was then assigned as the predicted class.

To benchmark the two strategies, we assessed the concordance between paired spatial and single-cell expression profiles. For each cell type, we subset both datasets to shared genes, log-normalised and scaled expression values within each dataset independently, and computed the mean expression profile per cell type. Concordance was quantified as the Pearson correlation between the average expression profile of cell type A in the single-cell data and the corresponding cell type A in the spatial data, across all cell types. On aggregate metrics, ISS-patcher without downsampling showed the highest overall concordance, while ISS-patcher with downsampling performed worst (**Extended Data Fig. 2b**). However, TACCO recovered more cell types overall (**Extended Data Fig. 2c**) and showed superior performance for rare cell types. For example, for the basalis  stromal fibroblasts introduced in Main Text and characterised by the expression of *SFRP5,* TACCO predictions most closely match the expression of *SFRP5* as measured in *Xenium* 480-plex (**Extended Data Fig. 2d**). Given the importance of accurately mapping rare populations, TACCO was used for all analyses in Main Text.

#### Spatial axis mapping onto scRNA-seq reference

To assign spatial positional information to cells in the scRNA-seq reference atlas, we transferred the myometrial-luminal axis (see Main Texts and Methods) by computationally pairing cells from the two modalities. To achieve this, we explored two complementary strategies. First, we applied ISS-patcher to assign each cell in the scRNA-seq data the average axis value of its k-nearest neighbors in spatial transcriptomics, thus directly transferring continuous axis coordinates. Second, we applied TACCO to here quantifying contributions of the six spatial compartments (basalis_1, basalis_2, functionalis_1, functionalis_2, functionalis_3, lumen_1, see Main Text) to each single cell transcriptome. For each single cell, the spatial compartment with the highest predicted contribution was assigned as its positional class, retaining only high-confidence assignments (maximum score > 0.4 for TACCO, axis std < 0.15 for ISS-patcher). Both methods were run with default parameters.

All transfer was performed separately for epithelial and stromal compartments, and restricted to stage-matched cells, whereby single-cell profiles were paired within the same menstrual phase. When multiple spatial samples were available for a given stage, we considered two strategies. In the first configuration ("average"), coordinate transfer was repeated independently with each spatial sample used as reference, then predicted scores were averaged across runs. In the second configuration ("concat"), the multiple spatial samples were first concatenated, then the pairing method was only run once.

To benchmark the two strategies, we assessed the concordance between paired spatial and single-cell expression profiles. For each lineage/phase combination, we subset both datasets to shared genes, log-normalised and scaled expression values within each dataset independently, and computed the mean expression profile per spatial compartment bin. Concordance was quantified as the Pearson correlation between the average expression profile of bin *i* in the single-cell data and the corresponding bin *i* in the spatial data, across all bins. TACCO "average" showed higher mean concordance (**Extended Data Fig. 4a**) and was used for all analyses in Main Text.

We then confirmed that paired axis compartments show higher pairwise gene expression correlation between single-cell and spatial data than non-paired compartments (**Extended Data Fig. 4b**). The resulting axis was further qualitatively validated using known spatial polarisation markers. Genes with previously reported basalis-biased expression (*CDH2, AXIN2, TRH, ALDH1A1, KLK11*)[3–5] were enriched in basal bins, while luminal markers *WNT7A* and *LGR5* were enriched in functionalis bins closest to the lumen, as assessed by visual inspection of dot plots of marker expression across positional bins. (**Extended Data Fig. 4c)**.

**Supplementary Note references**


1. To, K. *et al.* A multi-omic atlas of human embryonic skeletal development. *Nature* **635**, 657–667 (2024).
2. Mages, S. *et al.* TACCO unifies annotation transfer and decomposition of cell identities for single-cell and spatial omics. *Nat Biotechnol* **41**, 1465–1473 (2023).
3. Nguyen, H. P. T. *et al.* N-cadherin identifies human endometrial epithelial progenitor cells by in vitro stem cell assays. *Hum. Reprod.* **32**, 2254–2268 (2017).
4. Marečková, M. *et al.* An integrated single-cell reference atlas of the human endometrium. *Nat. Genet.* **56**, 1925–1937 (2024).
5. Fitzgerald, H. *et al.* Molecular signature of human endometrial stem/progenitor cells at the single cell level. *bioRxiv* 2025.06.23.660982 (2025) doi:10.1101/2025.06.23.660982.


#### Extended Data and Extended Data Figure legends

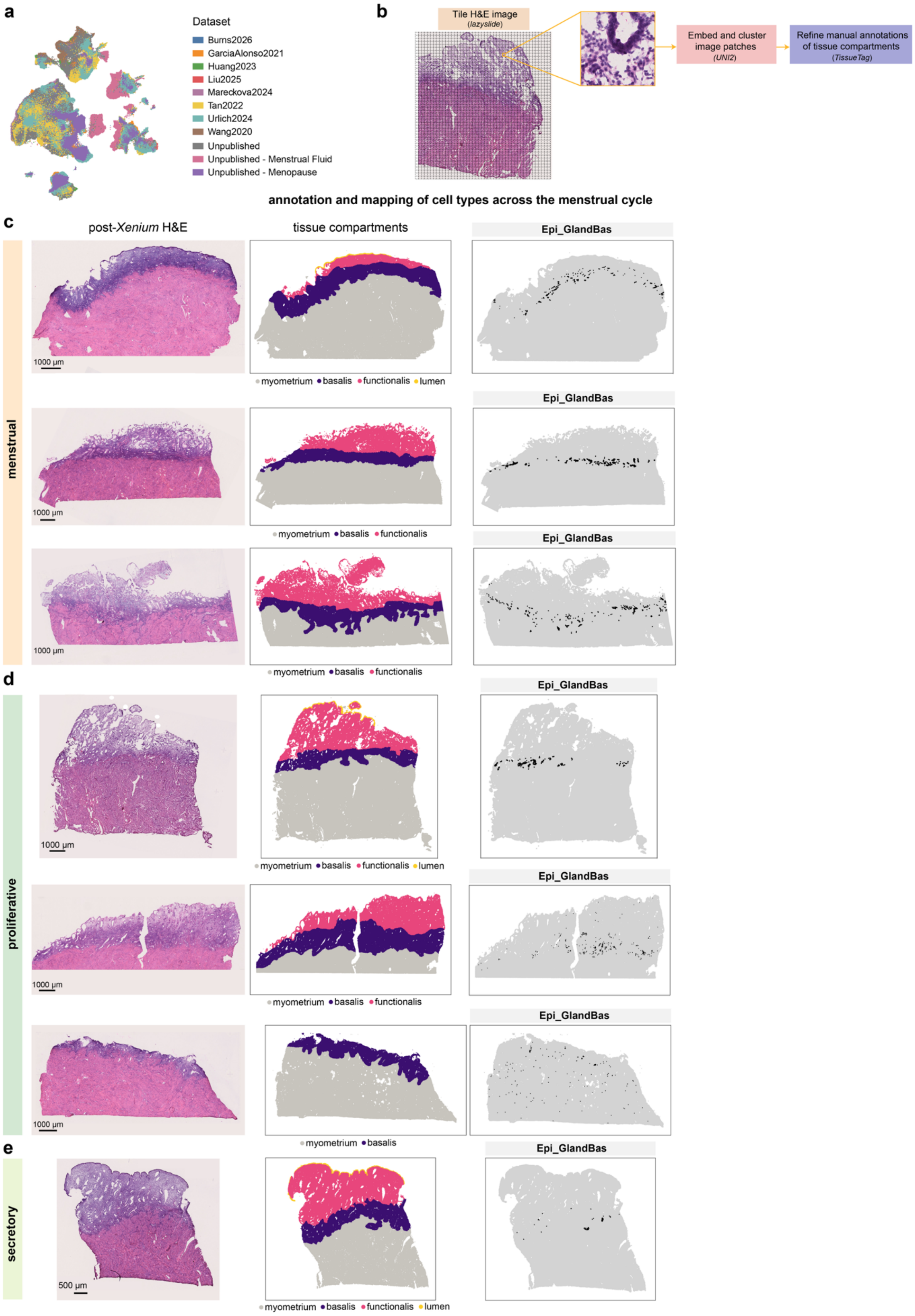
a
Dataset
Burns2026
GarciaAlonso2021
Huang2023
Liu2025
Mareckova2024
Tan2022
Urlich2024
Wang2020
Unpublished
Unpublished - Menstrual Fluid
Unpublished - Menopause
b
Tile H&E image
(lazyslide)
Embed and cluster image patches
(UNI2)
Refine manual annotations of tissue compartments
(TissueTag)
annotation and mapping of cell types across the menstrual cycle
c
post-Xenium H&E
tissue compartments
Epi_GlandBas
menstrual
1000 μm
myometrium basalis functionalis lumen
myometrium basalis functionalis
d
proliferative
myometrium basalis
e
secretory
500 μm

**Extended Data Figure 1**. **a,** UMAP embeddings of scRNA-seq cells in the Human Female Reproductive System Cell Atlas v1 coloured by dataset of origin. **b,** Schematic illustration of the computational workflow for annotating tissue compartments in post-*Xenium* H&Es. **c,** For each donor, post-*Xenium* H&E images of endometrial tissue sections (left) are shown alongside corresponding annotated tissue compartments (middle), including myometrium, basalis, functionalis, and lumen. The inferred spatial localisation of basalis epithelial glands (defined in scRNA-seq) is shown on the right. Donors are ordered by menstrual cycle phase.

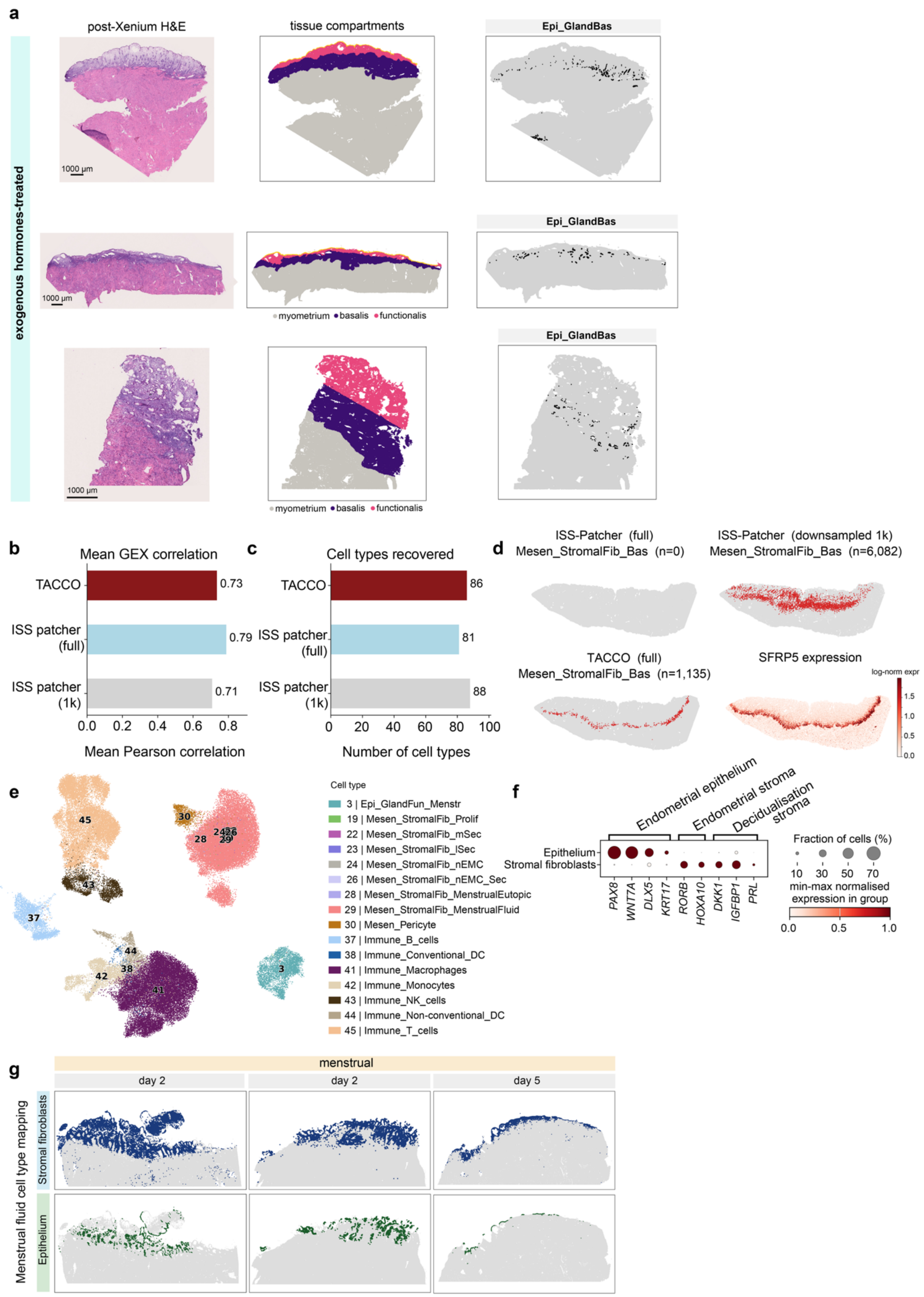


**Extended Data Figure 2. a,** As in **Extended Data Figure 1**, for each hormone-treated donor, post-*Xenium* H&E images of endometrial tissue sections (left) are shown alongside corresponding annotated tissue compartments (middle), including myometrium, basalis, functionalis, and lumen. The inferred spatial localisation of basalis epithelial glands (defined in scRNA-seq) is shown on the right. **b,** Barplot showing the mean Pearson correlation between the mean scaled gene expression of equivalent cell type classes in single cell and spatial transcriptomics, grouped by method. For ISS-patcher (full), the entire scRNA-seq dataset is used as a reference, for ISS-patcher 1k, fine cell types are downsampled to a maximum of 1000 cells per cell type. **c,** Average number of cell types recovered in spatial transcriptomics samples by each computational method. **d**, Predicted spatial localisations of the basalis stromal fibroblast population by ISS-Patcher (full; n=0 cell), ISS-Patcher (downsampled 1k; n=6,082 cells), and TACCO (full; n=1,135 cells), alongside spatial expression of the marker gene *SFRP5*, in a *Xenium* 480-plex spatial transcriptomics section from proliferative phase, demonstrating concordance between TACCO mapping and marker expression. **e,** UMAP embeddings of the menstrual fluid dataset colored by fine cell type. Annotations follow the numbering of **Figure 1b**. **f,** Dotplot showing the expression of canonical endometrial epithelial and stromal markers in epithelial and stromal cells from menstrual fluid. Dot size indicates the fraction of expressing cells, and colour intensity reflects min-max normalised expression within each group. **g,** Predicted spatial localisations of menstrual fluid epithelium and stromal fibroblasts on *Xenium* 480-plex spatial transcriptomics sections from menstrual days 2 (2 samples) and 5.

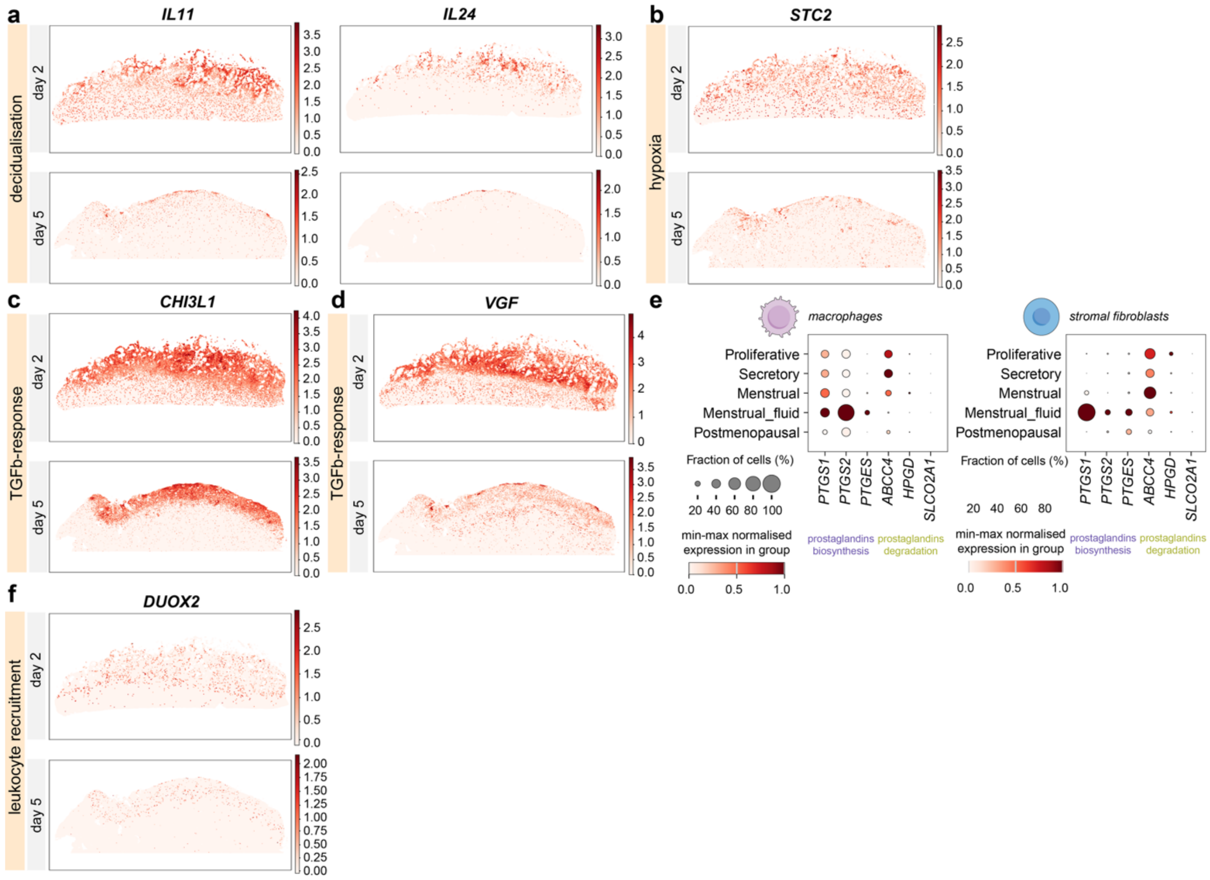


**Extended Data Figure 3. a,** Log-normalised expression of menstrual stromal markers related to decidualisation (*IL11*, *IL24*) in *Xenium* 5K+100-plex spatial transcriptomics sections from menstrual days 2 and 5. **b,** Log-normalised expression of the hypoxia stromal marker *STC2* in *Xenium* 5K+100-plex spatial transcriptomics sections from menstrual days 2 and 5. **c,** Log-normalised expression of the TGFβ-response stromal marker *CHI3L1* in

*Xenium* 5K+100-plex spatial transcriptomics sections from menstrual days 2 and 5. **d,** Log-normalised expression of the nociceptive stromal marker *VGF* in *Xenium* 5K+100-plex spatial transcriptomics sections from menstrual days 2 and 5. **e,** Dotplot showing the expression of prostaglandin biosynthesis and degradation genes in macrophages (left) and stromal fibroblasts (right) across menstrual phases. Dot size indicates the fraction of expressing cells, and colour intensity reflects min-max normalised expression within each group. **f,** Log-normalised expression of the immune recruitment epithelial marker *DUOX2* in *Xenium* 5K+100-plex spatial transcriptomics sections from menstrual days 2 and 5.

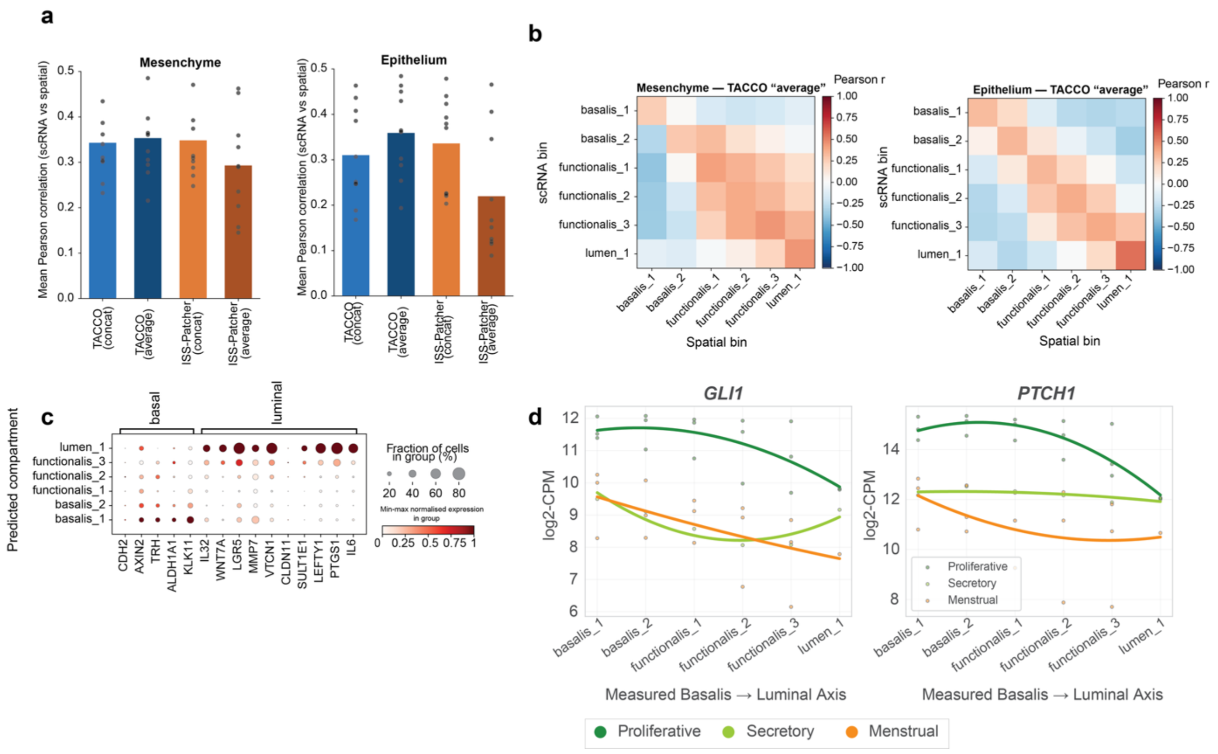


**Extended Data Figure 4. a,** Benchmarking spatial information transfer approaches. Barplot showing the mean Pearson correlation between matched spatial and single-cell compartment expression profiles following transfer of spatial information using TACCO or ISS-patcher, implemented using either concatenated spatial references ("concat") or averaged predictions across individual references ("average"). Points represent individual lineage-phase combinations. **b,** Heatmap showing the Pearson correlation between mean scaled gene expression profiles of scRNA-seq (rows) and spatial transcriptomic compartments (columns) across the basalis-to-luminal binned axis for mesenchymal (left) and epithelial (right) cells. Mappings are from the TACCO "average" configuration. **c,** Dot plot showing the expression of epithelial marker genes known to have basalis or luminal-biased expression across transferred basalis-to-luminal binned axis compartments in scRNA-seq data. Dot size indicates the fraction of expressing cells, and colour intensity reflects min-max normalised expression within each group. **d,** Spline plots showing the log-transformed expression of hedgehog signalling response genes in stromal fibroblast cells across the basalis-to-luminal binned axis in proliferative, secretory, and menstrual phases, as measured by *Xenium* 480-plex spatial transcriptomics. Lines are coloured by menstrual phase, and each point represents pseudobulked cells assigned to a specific bin per donor.

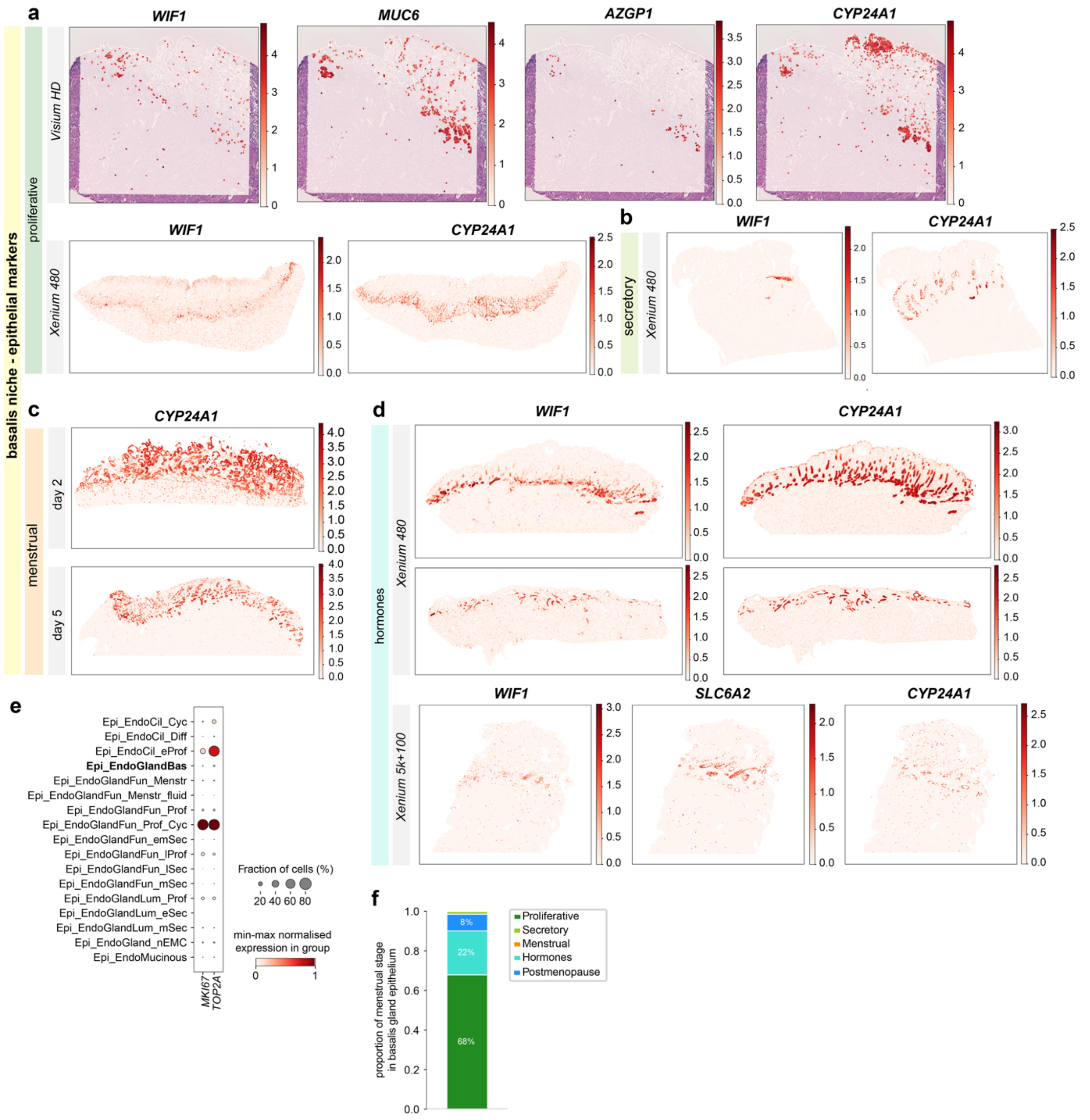


**Extended Data Figure 5. a,** Log-normalised spatial expression of basalis glands epithelial markers on a *Visium HD* spatial transcriptomics section from proliferative phase (top) and on a *Xenium* 480-plex spatial transcriptomics section from proliferative phase (bottom). **b,** Log-normalised spatial expression of basalis glands epithelial markers on a *Xenium* 480-plex spatial transcriptomics section from secretory phase. **c,** Log-normalised spatial expression of basalis glands epithelial markers on two *Xenium* 480-plex spatial transcriptomics sections from menstrual days 2 and 5. **d,** Log-normalised spatial expression of basalis glands epithelial markers on three *Xenium* 480-plex spatial transcriptomics sections (top) or *Xenium* 5k+100-plex (bottom) from donors taking exogenous hormones. **e,** Dotplot showing the expression of cell proliferation marker genes across epithelial cells in scRNA-seq data. Dot size indicates the fraction of expressing cells, and colour intensity reflects min-max normalised expression within each group. **f,** Barplot showing the proportion of basalis glandular epithelial cells colored by menstrual phase of the corresponding donor.

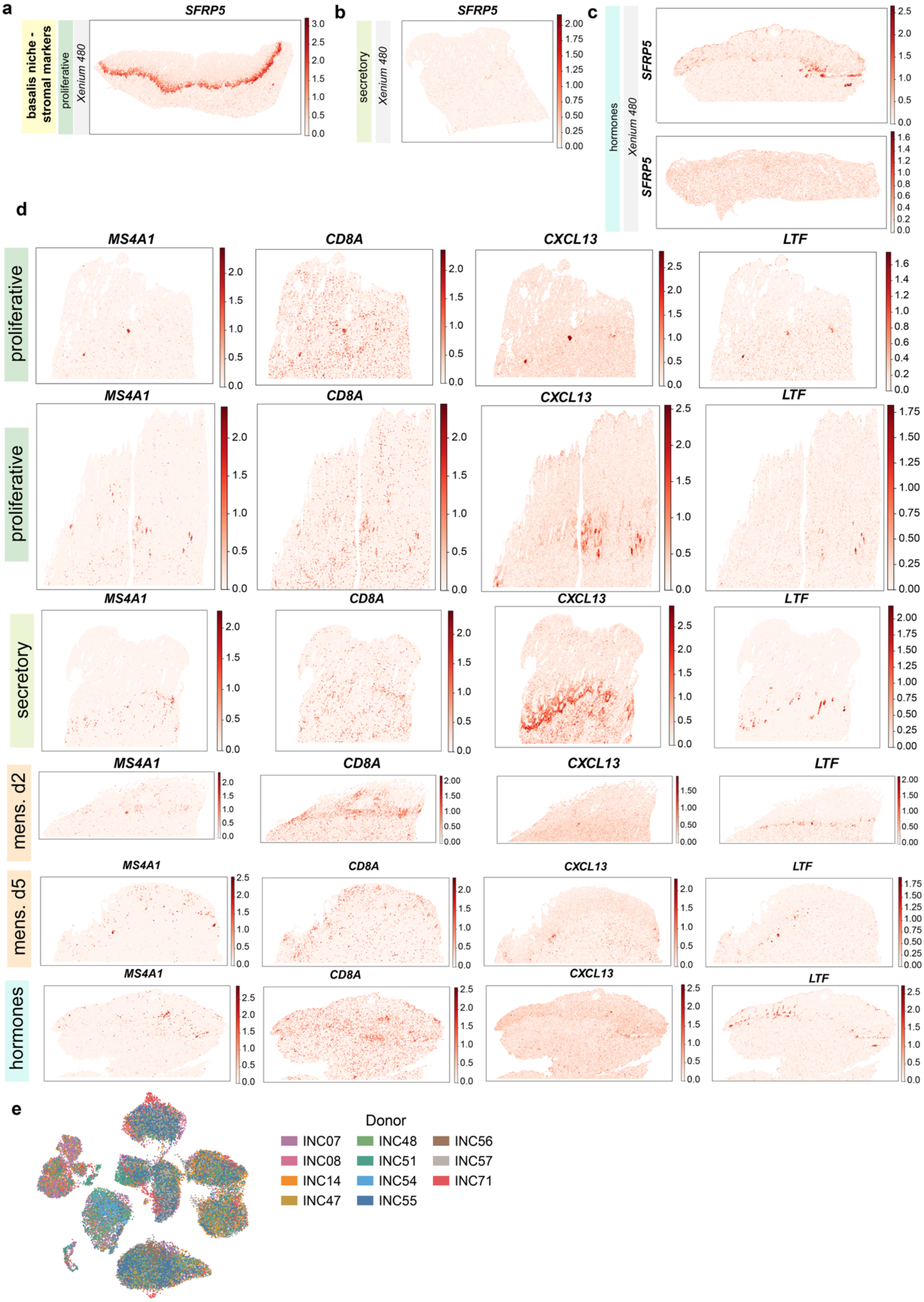

a
basalis niche - stromal markers
proliferative
Xenium 480
SFRP5
b
secretory
Xenium 480
SFRP5
c
hormones
Xenium 480
SFRP5
SFRP5
d
proliferative
proliferative
secretory
mens. d2
mens. d5
hormones
MS4A1
CD8A
CXCL13
LTF
e
Donor
INC07
INC08
INC14
INC47
INC48
INC51
INC54
INC55
INC56
INC57
INC71

**Extended Data Figure 6: a,** Log-normalised spatial expression of the basalis stromal fibroblast marker *SFRP5* on a *Xenium* 480-plex spatial transcriptomics section from proliferative phase. **b,** Log-normalised spatial expression of the basalis stromal fibroblast marker *SFRP5* on a *Xenium* 480-plex spatial transcriptomics section from secretory phase. **c,** Log-normalised spatial expression of the basalis stromal fibroblast marker *SFRP5* on *Xenium* 480-plex spatial transcriptomics sections from donors taking exogenous hormones. **d,** Log-normalised expression of lymphoid aggregate markers across donors. Each row corresponds to one donor, with menstrual phase indicated on the left hand side panel. **e,** UMAP embedding of the INCLIVA post-menopausal scRNA-seq dataset, coloured by donor.

#### Supplementary Table Legends

**Supplementary Table 1**: Patient metadata and sample-level quality control (QC) metrics for spatial transcriptomics samples, grouped by spatial transcriptomics panel and technology.

**Supplementary Table 2:** *Xenium* transcriptomic panel custom genes, grouped by panel.

**Supplementary Table 3**: Patient metadata sample-level QC metrics for newly generated scRNA-seq samples for the main Human Uterine Spatial Cell Atlas and for the INCLIVA post-menopausal validation dataset

**Supplementary Table 4:** Top-20 specific marker genes as obtained from TF-IDF by lineage (computed across the entire dataset) or by cell type (computed within each lineage).

**Supplementary Table 5:** Number of cells mapped to each spatial compartment for each cell type, by stage, grouped by lineage. For menstrual stages which include multiple tissue sections, values represent sums across donors.

**Supplementary Table 6**: Generalised linear model results, grouped by lineage. Empty rows for the interaction p-value correspond to genes for which the likelihood of the reduced model was greater than the likelihood of the interaction model (epithelial only). FDR_A corresponds to the adjusted p-value for the Wald-test for the axis coefficient. FDR_B corresponds to the adjusted p-value for the likelihood ratio test for the model with and without interaction term.
